# Annelid comparative genomics and the evolution of massive lineage-specific genome rearrangement in bilaterians

**DOI:** 10.1101/2024.05.15.594353

**Authors:** Thomas D. Lewin, Isabel Jiah-Yih Liao, Yi-Jyun Luo

## Abstract

The organization of genomes into chromosomes is critical for processes such as genetic recombination, environmental adaptation, and speciation. All animals with bilateral symmetry inherited a genome structure from their last common ancestor that has been highly conserved in some taxa but seemingly unconstrained in others. However, the evolutionary forces driving these differences and the processes by which they emerge have remained largely uncharacterized. Here we analyze genome organization across the phylum Annelida using 23 chromosome-level annelid genomes. We find that while most annelids have maintained the conserved bilaterian genome structure, a group containing leeches and earthworms possesses completely scrambled genomes. We develop a rearrangement index to quantify the extent of genome structure evolution and show leeches and earthworms to have the most highly rearranged genomes of any currently sampled bilaterian. We further show that bilaterian genomes can be classified into two distinct categories—high and low rearrangement—largely influenced by the presence or absence, respectively, of chromosome fission events. Our findings demonstrate that animal genome structure can be highly variable within a phylum and reveal that genome rearrangement can occur both in a gradual, stepwise fashion or as rapid, all-encompassing changes over short evolutionary timescales.

## Introduction

The arrangement of genomic DNA into individual chromosomes creates a dynamic landscape subject to continual reshaping through processes such as fusion, fission, inversion, and translocation (Schubert 2007; Lysák and Schubert 2013; Simakov et al. 2022). These structural alterations play pivotal roles in fundamental biological phenomena, including recombination (Rieseberg 2001; Näsvall et al. 2023; Yoshida et al. 2023), adaptation (Dunham et al. 2002; Coyle and Kroll 2007; Lowry and Willis 2010; Jones et al. 2012; Wellenreuther and Bernatchez 2018), speciation (Noor et al. 2001; de Vos et al. 2020; Augustijnen et al. 2024), disease (Lupski and Stankiewicz 2005), and ultimately, the emergence of novel phenotypic traits. While some lineages exhibit remarkable conservation of genome structure over extended evolutionary timescales, others display striking divergence, with chromosomes rearranged in unpredictable patterns (Wang et al. 2017; Simakov et al. 2022; Ivankovic et al. 2023; Lin et al. 2023; Martín-Zamora et al. 2023). Unraveling the evolutionary forces driving such lineage-specific scrambling of gene sets can provide valuable insights into the process of adaptation and the evolution of animal diversity.

The sequencing of genomes to chromosome scale has facilitated the development of new methods for comparing genome structure. In many species, orthologous genes have remained clustered on the same chromosomes for over half a billion years since the ancestor of bilaterian animals (Simakov et al. 2013; Simakov et al. 2022). This conserved gene linkage, or macrosynteny, can be used to track orthologous chromosomes across highly divergent species and is emerging as a powerful tool for studying genome rearrangements. This technique has thus far largely been applied to long-range comparisons across metazoans (Simakov et al. 2013; Simakov et al. 2022; Lin et al. 2023; Schultz et al. 2023; Zimmermann et al. 2023) or to compare very closely related species, for example *Acropora* corals (Locatelli et al. 2023) or cryptic species of the tunicate *Oikopleura dioica* (Plessy et al. 2024). Such studies have been highly fruitful, elucidating the genome structure of the last common ancestor of bilaterians (Putnam et al. 2007; Putnam et al. 2008; Simakov et al. 2022; Marlétaz et al. 2023) and developing algorithms for inferring rearrangement history (Ferretti et al. 1996; DasGupta et al. 1997; Mackintosh et al. 2023). There is, however, a lack of understanding about how genome structure evolves at a resolution between these two extremes, at the level of individual phyla. Key questions persist, including the degree of macrosynteny conservation within phyla, the frequency of large-scale reorganization events at lower taxonomic levels, the relative significance of fusion versus fission events, and how rearrangements unfold—whether gradually and stepwise or through sweeping changes over relatively short timescales.

The phylum Annelida represents a promising model to answer such questions. At the onset of this project, 24 chromosome-level annelid genomes were available from 15 families, including the basal owneniids, offering a dataset with both depth and breadth of sampling. Annelids form a diverse group of segmented, vermiform (worm-like) spiralians that is split into two main clades, Errantia and Sedentaria, based on the dominant lifestyle of their members (i.e. errant or sedentary) (Bleidorn et al. 2015). Annelid species such as the bristle worm *Capitella teleta*, the ragworm *Platynereis dumerilii* and the leech *Helobdella robusta* have emerged as key model systems, especially for questions surrounding the evolution of development (Weisblat and Kuo 2009; Kutschera and Weisblat 2015; Seaver 2016; Özpolat et al. 2021). While annelids are ancestrally ocean-dwelling and the majority of annelid diversity remains marine, the subclass Clitellata, containing earthworms and leeches, made a highly successful foray into freshwater and terrestrial habitats (Bleidorn et al. 2015; Erséus et al. 2020). Notably, it has been reported from draft assemblies that conserved bilaterian chromosomes are present in *C. teleta* and the miniature annelid *Dimorphilus gyrociliatus* but not in the leech *H. robusta* or the earthworm *Eisenia andrei*, pointing towards the possibility of extensive chromosome rearrangements within this group (Simakov et al. 2013; Martín-Durán et al. 2021; Sun et al. 2021).

Here, we first produce gene annotations for 22 chromosome-level annelid genomes. Using these new gene models, we build an updated phylogeny of annelids and characterize annelid chromosome evolution. Our findings suggest that the last annelid common ancestor had a genome of 20 chromosomes, with four fusion events compared to the ancestor of bilaterians. This karyotype is deviated from only slightly within most extant annelids, though fusion events are relatively frequent and have resulted in a chromosome number of less than 20 in all except one of the analyzed species. However, this conserved genome architecture has completely disintegrated within leeches and earthworms, and genes from conserved bilaterian linkage groups are shuffled across chromosomes. Using a newly defined rearrangement index, a metric aimed at quantifying the extent of chromosome rearrangement within genomes, we show that leeches and earthworms have among the highest levels of genome rearrangements of any bilaterian.

## Results

### Phylogeny of chromosome-level annelid genomes

To study annelid genome evolution, we first assembled a dataset of all public chromosome-level annelid genomes (supplementary table S1) and performed gene prediction using available RNA sequencing data. This produced a dataset of 23 chromosome-level assemblies with highly complete gene models (supplementary fig. S1). Genome size is highly variable among the sampled annelids, ranging from 149 Mb in the leech *Hirudinaria manillensis* to 1,861 Mb in the deep-sea hydrothermal vent scale worm *Branchipolynoe longqiensis* (mean assembly length 944 Mb) (supplementary fig. S2). Across the sampled genomes, the mean GC content is 39.6% and the mean repeat content is 45.8%. The wide range of chromosome numbers, from 9 to 41 with a mean of 16, makes this dataset particularly promising for studying interchromosomal rearrangements.

Robust phylogenies form the foundations for understanding the direction of evolutionary change and are therefore a necessity for studying genome evolution. However, the current understanding of annelid phylogeny is largely based on transcriptomic data (Struck et al. 2011; Weigert et al. 2014; Andrade et al. 2015; Weigert and Bleidorn 2016) and subsequently retains a degree of uncertainty. We built a maximum likelihood phylogeny of annelids using the chromosome-level genomes and newly annotated gene models (fig. 1; supplementary fig. S3). The topology is largely consistent with transcriptome-based phylogenies and supports the widely accepted division of the bulk of annelid diversity into two monophyletic groups, Errantia and Sedentaria, with Oweniidae and Sipuncula as basal lineages. Clitellata, including leeches and earthworms, form a clade that is sister to a clade containing Echiuroidea and Terebellida. This sister relationship of marine and freshwater clades highlights the lineage-specific evolution in habitat adaptation within the annelids.

**Fig. 1.**
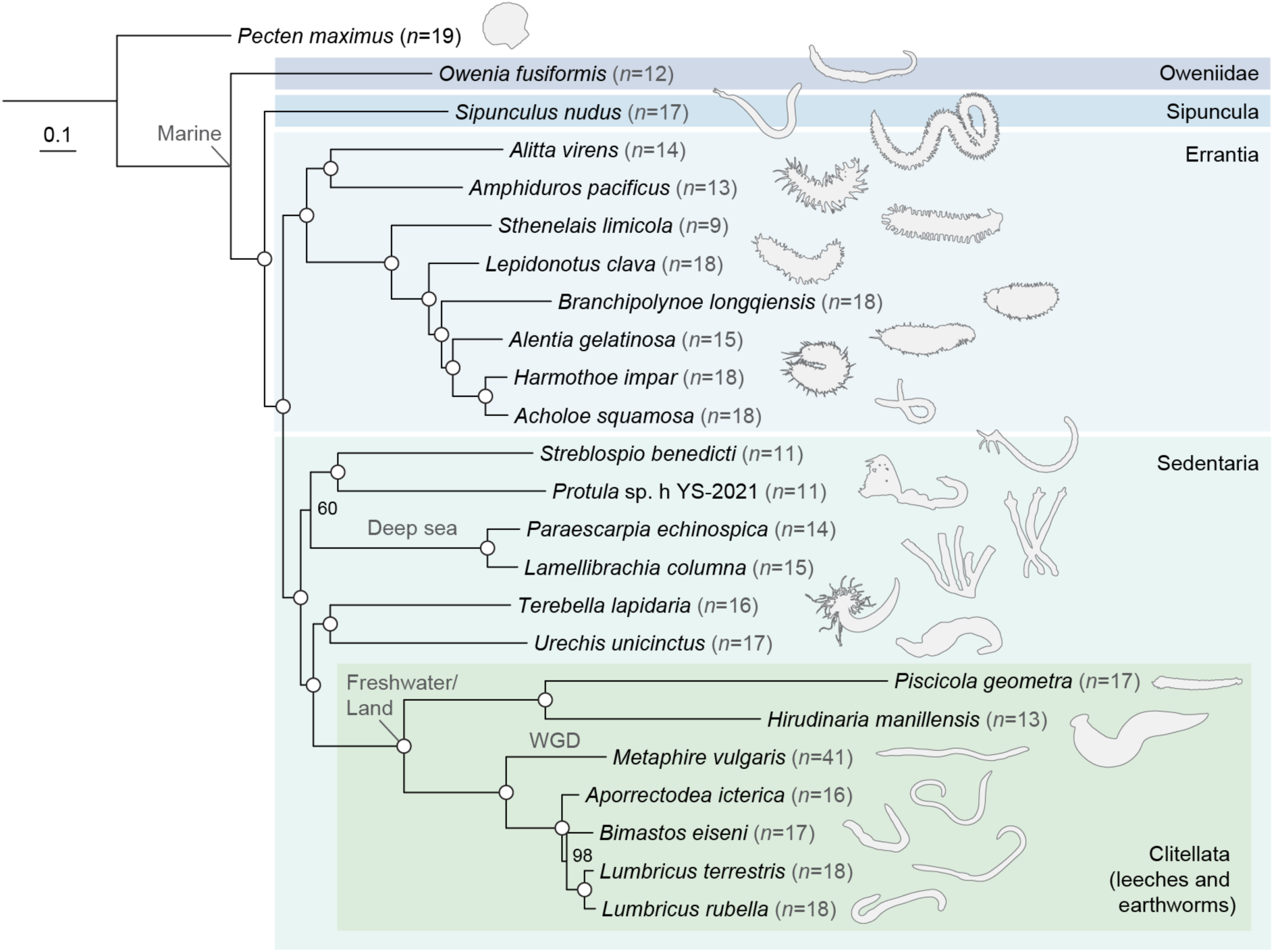
Phylogenomic analysis of annelids with chromosome-level genomes highlights lineage-specific evolution. Phylogenetic tree was constructed using the maximum likelihood method with the LG+F+R7 model based on a concatenated alignment of 537 single-copy orthologs from 23 annelid genomes. The scallop *P. maximus* is used as an outgroup. Open circles represent a bootstrap score of 100% support. Numbers in parentheses show chromosome number. WGD, whole genome duplication.

### Bilaterian ancestral linkage groups are often fused but rarely split in annelids

We next employed a macrosynteny approach to study interchromosomal rearrangements in annelids. To achieve this, we identified single-copy orthologs, assigned them to bilaterian ancestral linkage groups (ALGs) based on protein sequence homology, mapped their genomic locations, and used ideogram plots (fig. 2) and Oxford dot plots (supplementary figs. S4 and S5) to track chromosome relationships between species (supplementary table S2). The last bilaterian common ancestor had 24 ALGs, sets of genes that were subsequently inherited by all bilaterian phyla (Simakov et al. 2022), and we first questioned whether these are conserved in annelids. We found that all 24 ALGs were present in annelids and that there are four rearrangements to this structure that are shared by all annelids: H⊗Q, J2⊗L, K⊗O2, and O1⊗R (where ⊗ represents ALG fusion followed by gene mixing). From this, it can be inferred that the annelid ancestral state was 20 ALGs and, therefore, 20 chromosomes. Indeed, these four rearrangements are also shared by other lophotrochozoans, including molluscs, nemerteans, bryozoans, and brachiopods (Simakov et al. 2022; Lewin et al. 2024), suggesting they are common to most or all Lophotrochozoa. Therefore, there are no unique genome rearrangements shared by all annelids, and the last annelid common ancestor retained the ancestral lophotrochozoan karyotype with 20 chromosomes. Further, there are no chromosome rearrangements that act as synapomorphies for either of the large annelid clades, Errantia or Sedentaria, both of which inherited the 20 chromosomes possessed by the last common ancestor of annelids (fig. 2; supplementary fig. S6).

**Fig. 2.**
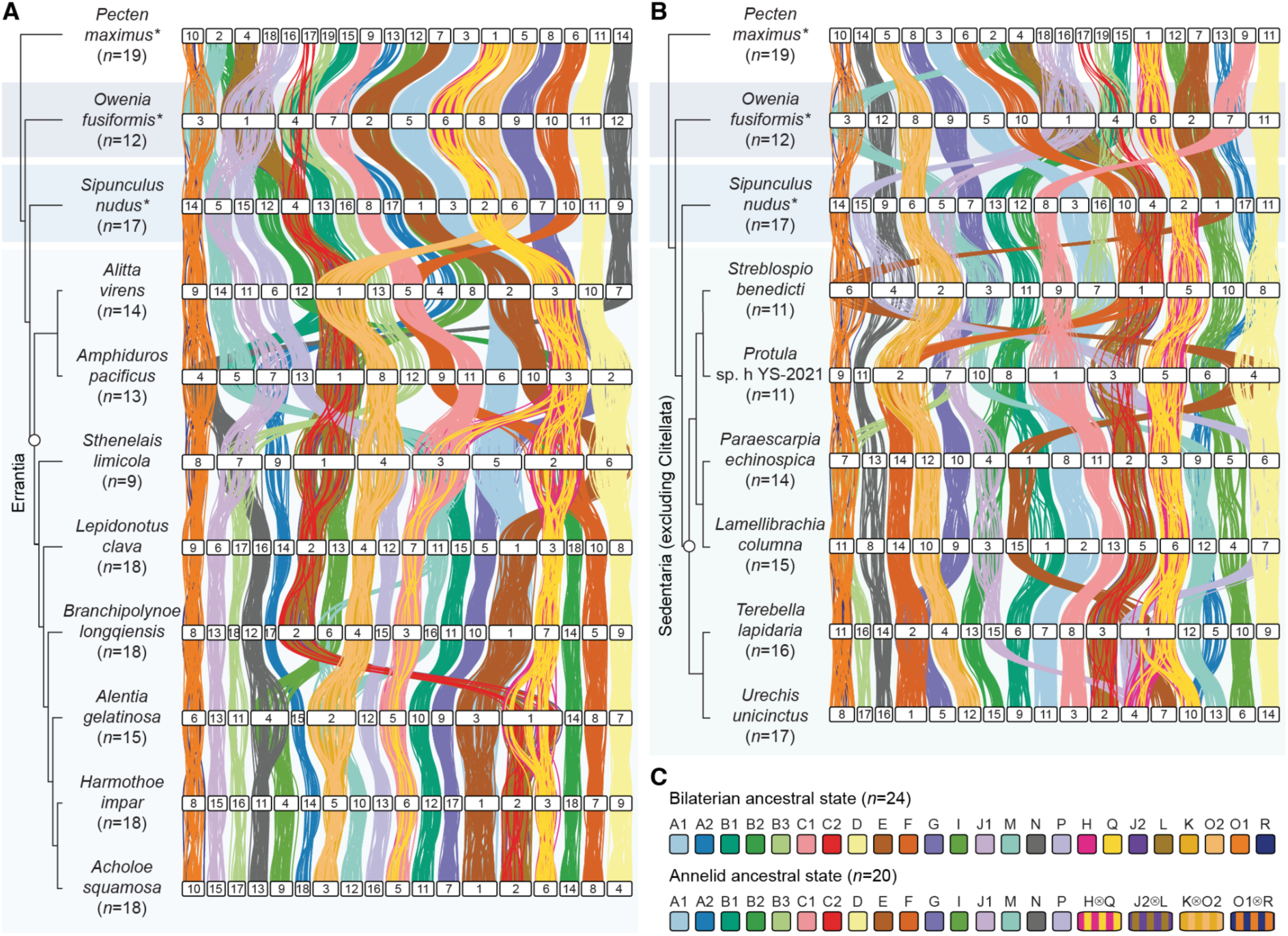
Bilaterian ancestral linkage groups are often fused but rarely split in errantian and sedentarian annelids. Ideogram plots displaying the locations of shared single-copy orthologs on chromosomes in Errantia (**A**) and Sedentaria (**B**) annelids. Phylogenies show the relationships determined in fig. 1, with the clades Errantia and Sedentaria highlighted with open circles. The scallop *P. maximus*, together with the annelids *Owenia fusiformis* and *Sipunculus nudus*, are used as outgroups and are labeled with asterisks. Horizontal white bars represent chromosomes. Vertical lines link orthologous genes between species; lines are colored by the bilaterian ancestral linkage groups to which genes belong. (**C**) The bilaterian ancestral state consisted of 24 linkage groups, while the annelid ancestral state had 20 linkage groups, with the following fusions compared to the bilaterian ancestral state: H⊗Q, J2⊗L, K⊗O2, and O1⊗R. The symbol ⊗ represents fusion-with-mixing events.

In addition to these four chromosome fusion events present in all annelids, there are many further chromosome rearrangements restricted to specific lineages. The genomes of all sampled species have at least three ALG fusions compared to the ancestral annelid genome. There is only one case of two species sharing an identical genome structure in this dataset, *Harmothoe impar* and *Acholoe squamosa* (*n* = 18), which are both within the same family (Polynoidae). *Sthenelais limicola* (*n* = 9) has the highest number of fusions (i.e. 12) and *Sipunculus nudus* (*n* = 17), *Lepidonotus clava* (*n* = 18), *H. impar* (*n* = 18) and *A. squamosa* (*n* = 18) have the fewest (i.e. 3). Despite these differences, the distribution of fusion events between species does not vary significantly from that which would be expected by chance (chi-squared test, *P* = 0.214, df = 15) (supplementary fig. S7), suggesting that there is no variation in species’ propensity for chromosome fusions. We find that almost all the chromosome fusion events in annelids can be categorized as fusion-with-mixing, where ALG fusion is followed by gene shuffling. There are only three putative fusion events without mixing (notation ●): B1●E on *Paraescarpia echinospica* chromosome 1; (H⊗Q)●(E⊗P) on *Terebella lapidaria* chromosome 1; and G●(B3⊗J1) on *Protula sp. h YS-2021* chromosome 7. All other events are followed by extensive gene mixing.

A recent study in Lepidoptera (butterflies and moths) found that ALGs’ propensity for fusion was inversely correlated with the length of the chromosomes on which they reside (Wright et al. 2024). In annelids, we found no correlation between the ALG fusion rate and the length of chromosomes (Spearman’s rank correlation ρ(19) = 0.125, *P* = 0.607) (supplementary fig. S8, supplementary tables S3 and S4). Indeed, there was no significant difference in the rates at which ALGs fused (chi-squared test, *P* = 0.736, df = 18), suggesting that, in this annelid dataset, certain ALGs are not more prone to fusing than others.

In contrast to fusions, the splitting of ALGs is relatively rare, with only three cases in this dataset of 16 annelid species. The ALG H⊗Q is split independently in the suborder Aphroditiformia and in *Urechis unicinctus*, and the ALG M is split in *B. longqiensis*. We noted that ALG H⊗Q is housed on the second-longest chromosomes in the dataset, but more data is needed to determine whether ALG splitting is associated with chromosome length. In all cases, ALG splitting coincides with the fusion of part of an ALG to another chromosome rather than simple chromosome fission. Overall, ALG fusion is very common, but ALG splitting is rare within this dataset of annelid species.

### Total loss of bilaterian genome structure in leeches and earthworms

Annelid chromosome evolution is characterized by the broad maintenance of bilaterian ALGs with relatively frequent lineage-specific fusion events. Clitellata, including earthworms and leeches, is a morphologically divergent monophyletic group nested within the Sedentaria. Performing macrosynteny analysis on the genomes of six clitellates, we found that bilaterian ALGs have been completely lost in this clade (fig. 3A). Remarkably, synteny dot plots reveal that in both leeches and earthworms, there is complete shuffling of the ancient bilaterian genome, with each ALG spread across all chromosomes (fig. 3B). Interestingly, while genome structure is largely conserved within the leeches and earthworms, there has also been massive genome shuffling between these two groups. Their genomes cannot be easily mapped to each other and have highly divergent organizational structures (fig. 3B). Overall, the ancient bilaterian genome architecture has been completely lost within a specific clade of annelids, the clitellates.

**Fig. 3.**
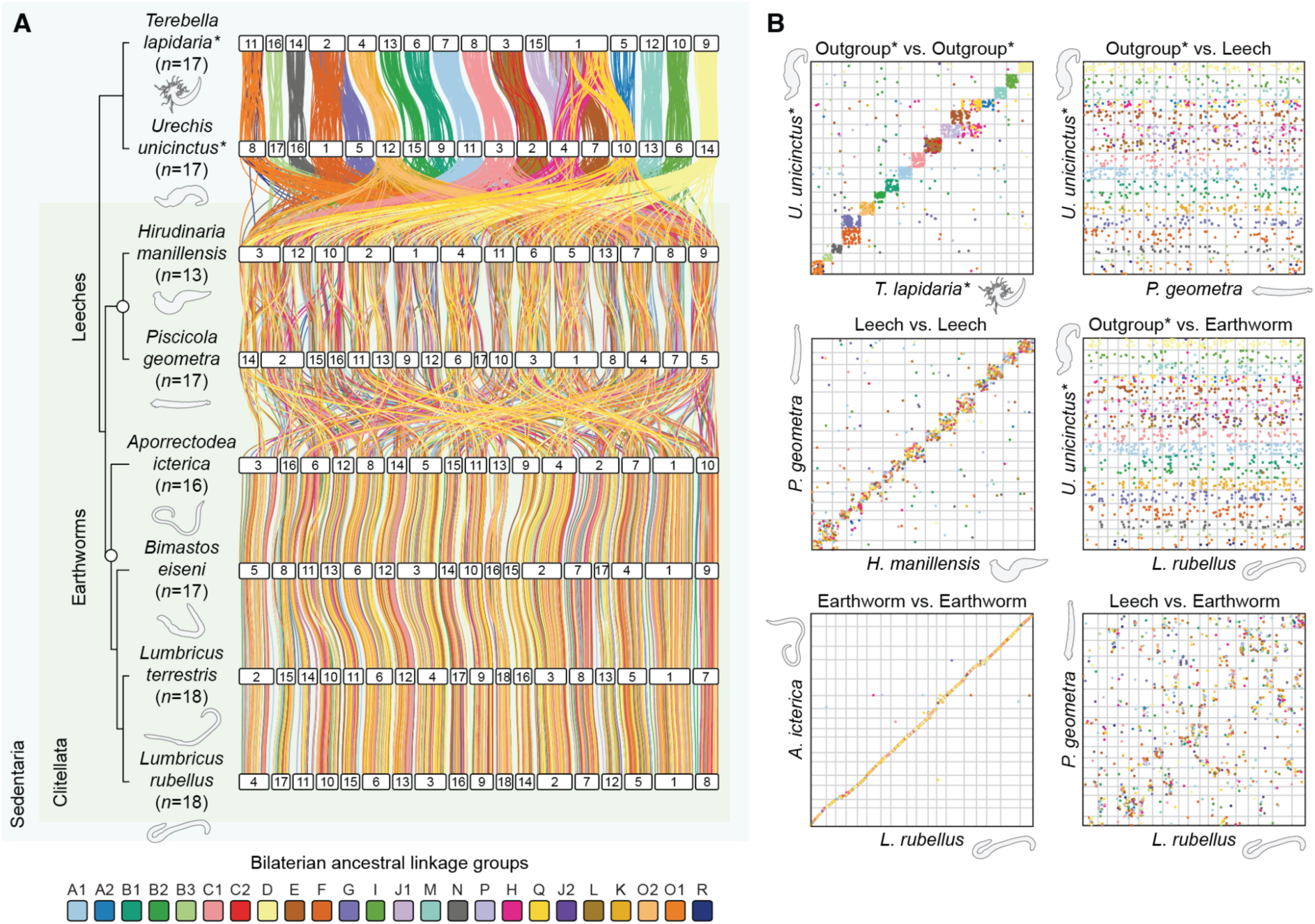
Bilaterian ancestral linkage groups are completely rearranged in leech and earthworm genomes. (**A**) Syntenic analysis of clitellate annelid genomes. Horizontal white bars represent chromosomes. Vertical lines connect orthologous genes between species and are colored according to the bilaterian ancestral linkage groups to which the genes belong. Phylogeny shows the topology determined in fig. 1. The non-clitellate annelids *T. lapidaria* and *U. unicinctus* are utilized as outgroups and are marked by asterisks. (**B**) Oxford dot plots of shared orthologous genes across chromosomes between pairwise annelid genomes. Dots organized into quadrangles indicate the conservation of macrosynteny (genes on the same chromosome) but not microsynteny (gene order on the chromosome), as seen in comparisons within the outgroup. In contrast, dots aligned in a straight line represent the conservation of both macrosynteny and microsynteny, such as in earthworm vs. earthworm comparisons. Dots scattered randomly across the plot without any clear organization suggest no conservation of macrosynteny or microsynteny.

### Lineage-specific whole genome duplication in earthworms

Whole genome duplication (WGD) is important to the evolution of morphological complexity and the emergence of evolutionary novelties (Moriyama and Koshiba-Takeuchi 2018). It has been reported that the genome of the earthworm *Metaphire vulgaris* underwent a WGD event leading to tetraploidization (Jin et al. 2020). Given the massive genome shuffling present in all clitellates, we questioned whether this genome duplication is specific to *M. vulgaris* or is present across all leeches and earthworms. Our bilaterian ALG-based macrosynteny approach suggests that the duplication is unique to *M. vulgaris*: we found that macrosynteny is partially conserved with other earthworms like *Aporrectodea icterica* but there is a 1:2 correspondence between many sections of the *A. icterica* genome with that of *M. vulgaris*. This 1:2 ratio is visible both in an ideogram plot (fig. 4A) and Oxford dot plot (fig. 4B; supplementary figs. S9 and S10) and is highly suggestive of a WGD in *M. vulgaris* but not *A. icterica*.

**Fig. 4.**
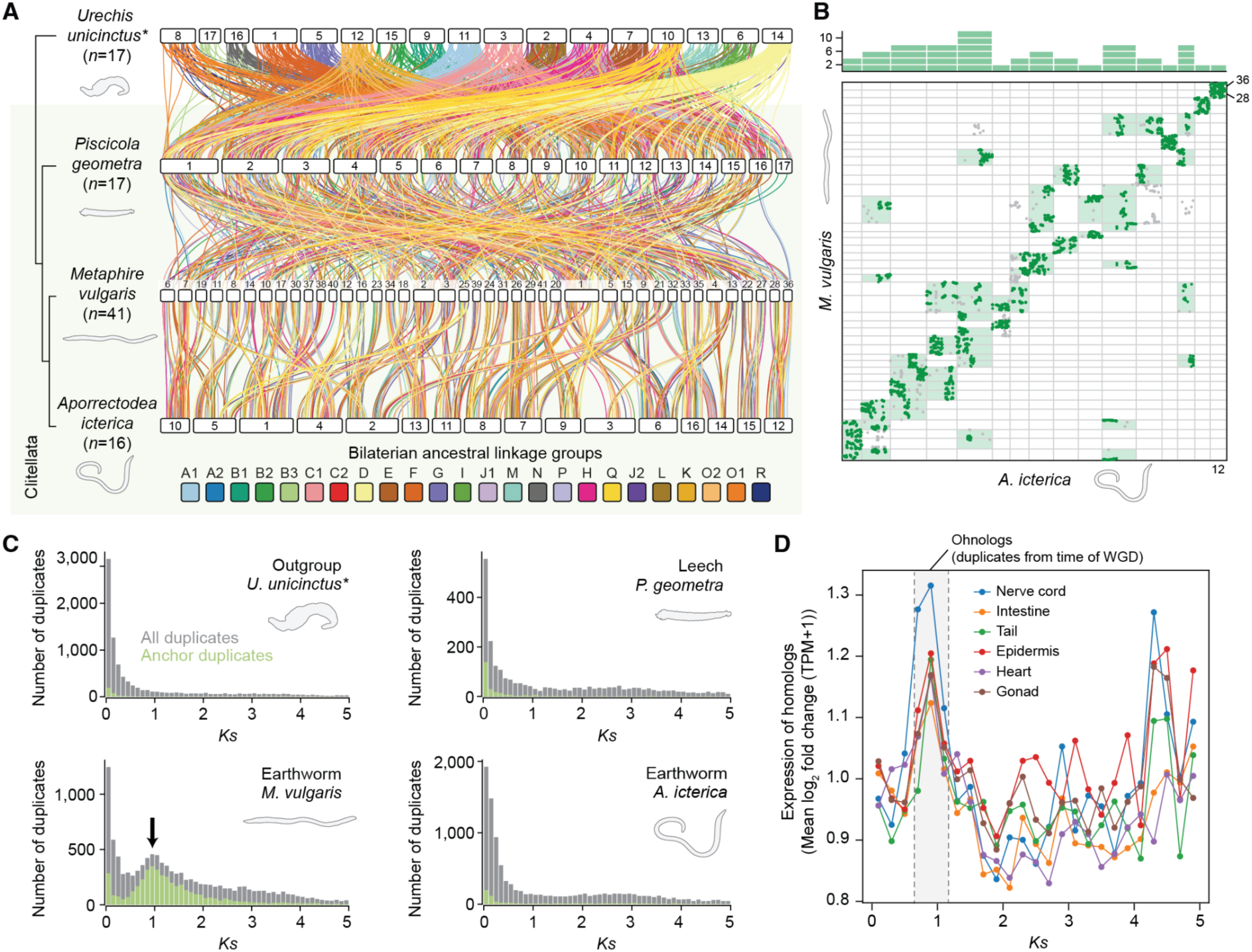
Organization of bilaterian ancestral linkage groups supports lineage-specific whole genome duplication in the earthworm *M. vulgaris*. (**A**) Ideogram plot of *M. vulgaris* (earthworm) genome with those of *A. icterica* (earthworm), *P. geometra* (leech) and *U. unicinctus* (outgroup). Horizontal bars represent chromosomes. Vertical lines link orthologous genes between species; lines are colored by the ALG to which genes belong. Phylogeny shows the topology determined in fig. 1. The *M. vulgaris* genome is completely rearranged compared to Sedentaria species and leeches but shows some conserved macrosynteny with other earthworms. Many areas of the earthworm *A. icterica* genome correspond to two areas of the *M. vulgaris* genome. Links between chromosomes with fewer than 20 orthologous genes are trimmed from the plot for clarity. (**B**) Oxford dot plot of *M. vulgaris* and *A. icterica* genomes, where each point represents an orthologous gene’s position in both genomes. Plot shows single-copy orthologs and one-to-many orthologs with up to five copies in one species. Sets of orthologues with a 1:2 ratio in the *A. icterica* versus *M. vulgaris* are shown in dark green. For instance, *A. icterica* chromosome 12 corresponds to *M. vulgaris* chromosomes 28 and 36. Boxes representing the chromosomes on which they appear are highlighted in light green. The bar chart at the top shows the number of cases of this on each *A. icterica* chromosome. The plot shows that many chromosome sections in *A. icterica* correspond to two chromosome sections in *M. vulgaris*, suggestive of whole genome duplication. A larger version of this figure with chromosomes labeled and duplicated regions highlighted is presented as supplementary fig. S10. (**C**) *K_S_* plots for duplicated gene families. Gray bars show histograms for all duplicated genes, while green bars show only anchor duplicates: these are duplicated genes found in duplicated collinear blocks of genes in the genome. Genomes with no whole genome duplication are expected to show exponential decay of duplicate numbers as *K_S_* increases, as seen in the *A. icterica* (earthworm), *P. geometra* (leech) and *U. unicinctus* (outgroup) plots. The *M. vulgaris* plot is interrupted by a normally distributed peak, indicative of whole genome duplication. (**D**) Difference of expression of *M. vulgaris* gene duplicate pairs (measured in log_2_ fold change of transcripts per million (TPM) + 1) in six tissues plotted against *K_S_*, a proxy for time since duplication. There is a peak of expression difference in gene duplicate pairs which emerged at the time of the whole genome duplication in all six tissues.

To test this, we first produced synonymous substitution rate (*K_S_*) plots for gene duplicates within each genome. *K_S_* measures the number of synonymous substitutions per site and, assuming that such changes are neutral and occur at a constant rate, estimates the evolutionary time since a gene duplication (Maere et al. 2005). In typical genomes with no WGD, the largest number of retained duplicates are evolutionarily young with low *K_S_*, and the number of retained duplicates decreases exponentially as *K_S_* increases. However, a WGD produces many duplicates at the same time, resulting in an excess of gene families of the same age; this manifests as a normally distributed peak at an intermediate *Ks* value (Blanc and Wolfe 2004; Schlueter et al. 2004; Tiley et al. 2018). We find exactly such a peak in the *K_S_* plot for *M. vulgaris* but not for other leeches, earthworms, or more distantly related annelids (fig. 4C; supplementary fig S11). This strongly supports the above conclusion of a recent WGD in *M. vulgaris*. Furthermore, the pattern is also repeated when only anchor gene duplicates are considered. Anchors are genes that are present in duplicated collinear blocks in the genome (i.e. conserved microsyntenic blocks), suggesting that they are not tandem duplicates and more likely arose from WGD events. This makes them more reliable for constructing *K_S_* plots; the presence of an anchor gene *K_S_* peak in *M. vulgaris* but not other species therefore further supports a specific WGD in this lineage (fig. 4C; supplementary fig S11).

Finally, we supplemented this data with structural information by searching for large collinear blocks of homologous genes within the annelid genomes. Most annelids do not have such blocks, but there are many pairs of locations in the *M. vulgaris genome* that share conserved microsynteny (supplementary figs. S12 and S13). This supports the presence of a recent WGD in *M. vulgaris* but not other annelids. Taken together, these data strongly support the conclusion that within the Clitellata, *M. vulgaris* has a lineage-specific tetraploidization that is not shared by other earthworms or leeches. Denser sampling is needed to determine whether the WGD is shared with other species within the family Megascolecidae. Interestingly, our result demonstrates that a macrosynteny approach using single-copy orthologs can reveal the presence of a WGD which is then supported by traditional detection methods such as analysis of synonymous substitution rate.

A key question emerging from these results is what biological effects these changes might have. We questioned whether gene duplicates formed by the WGD evolve differently to those formed by non-WGD duplication events. To answer this, we plotted duplicate pairs’ *Ks* values against the log_2_ fold difference in their expression level in RNA-sequencing datasets from six *M. vulgaris* tissues (supplementary tables S5 and S6). We found a peak in the difference of expression in duplicate pairs that emerged at the time of the WGD (fig. 4D). This suggests that duplicate genes derived from this WGD diverged in expression more quickly than those formed by other processes, such as tandem duplication, underlining the potential importance of WGD events to evolvability and adaptation.

### Exceptional levels of genome reorganization in clitellates

Next, we investigated whether the level of genome rearrangements observed in clitellates is unusual within the context of other bilaterians. To test this, we developed a macrosynteny rearrangement index, *R_i_* (see Materials and Methods; supplementary fig. S14 for a complete explanation), a metric to quantify both ALG fusion and fission into a single value between 0 (no rearrangement) and 1 (maximum rearrangement). The index considers macrosynteny (chromosomal colocalization) alone and does not consider microsynteny (conserved gene order). Unlike a previous conservation index (Simakov et al. 2013), it does not require pairwise comparisons and can therefore be computed for a single genome, and does not rest on potentially unreliable homology of chromosomes or scaffolds. The rearrangement index comprises an ALG splitting parameter and an ALG combining parameter. Though we expect the combining parameter to largely reflect ALG fusion and the splitting parameter to largely reflect ALG fission, the index cannot distinguish fusion and fission from other mechanisms, such as reciprocal translocation, so in the below discussion, we avoid the terms fusion and fission and instead use splitting (genes from one ALG being separated on to different chromosomes) and combining (genes from different ALGs coming together on the same chromosome.

We used the rearrangement index to compare rearrangement levels in our annelid dataset with species from 11 other bilaterian phyla for which chromosome-level assemblies were available (fig. 5A; supplementary fig. S15, supplementary table S7). We first validated our approach by comparing species previously reported to have highly rearranged genomes. The high rearrangement indices of octopus (*O. bimaculoides*, *n* = 30) (Albertin et al. 2022), diptera (flies) (*Drosophila melanogaster*, *n* = 4) (Wang et al. 2017), bryozoans (*C. mucedo*, *n* = 8) (Lewin et al. 2024) and platyhelminthes (*S. mansoni*, *n* = 11) (Wang et al. 2017; Ivankovic et al. 2023) demonstrate that the index correctly scores these species as highly rearranged. Conversely, deuterostomes such as starfish (*Asterias rubens*, *n* = 22) and sea cucumber (*Holothuria leucospilota*, *n* = 23) have the lowest levels of rearrangement, maintaining genome structures most similar to that of the bilaterian ancestor. The scallop (*P. maximus*, *n* = 19) and amphioxus (*B. floridae*, *n* = 19) also have highly conserved genomes with minimal rearrangement. Other species that we identify with highly rearranged genomes include fast-evolving lineages with few chromosomes like rotifers (*Adineta vaga*, *n* = 6) and parasites like nematomorphs (*Gordionus* sp. RMFG-2023, *n* = 5). Strikingly, clitellate annelids have the highest rearrangement index of all sampled bilaterians, suggesting that genomes in this group have been subject to the most rearrangement.

**Fig. 5.**
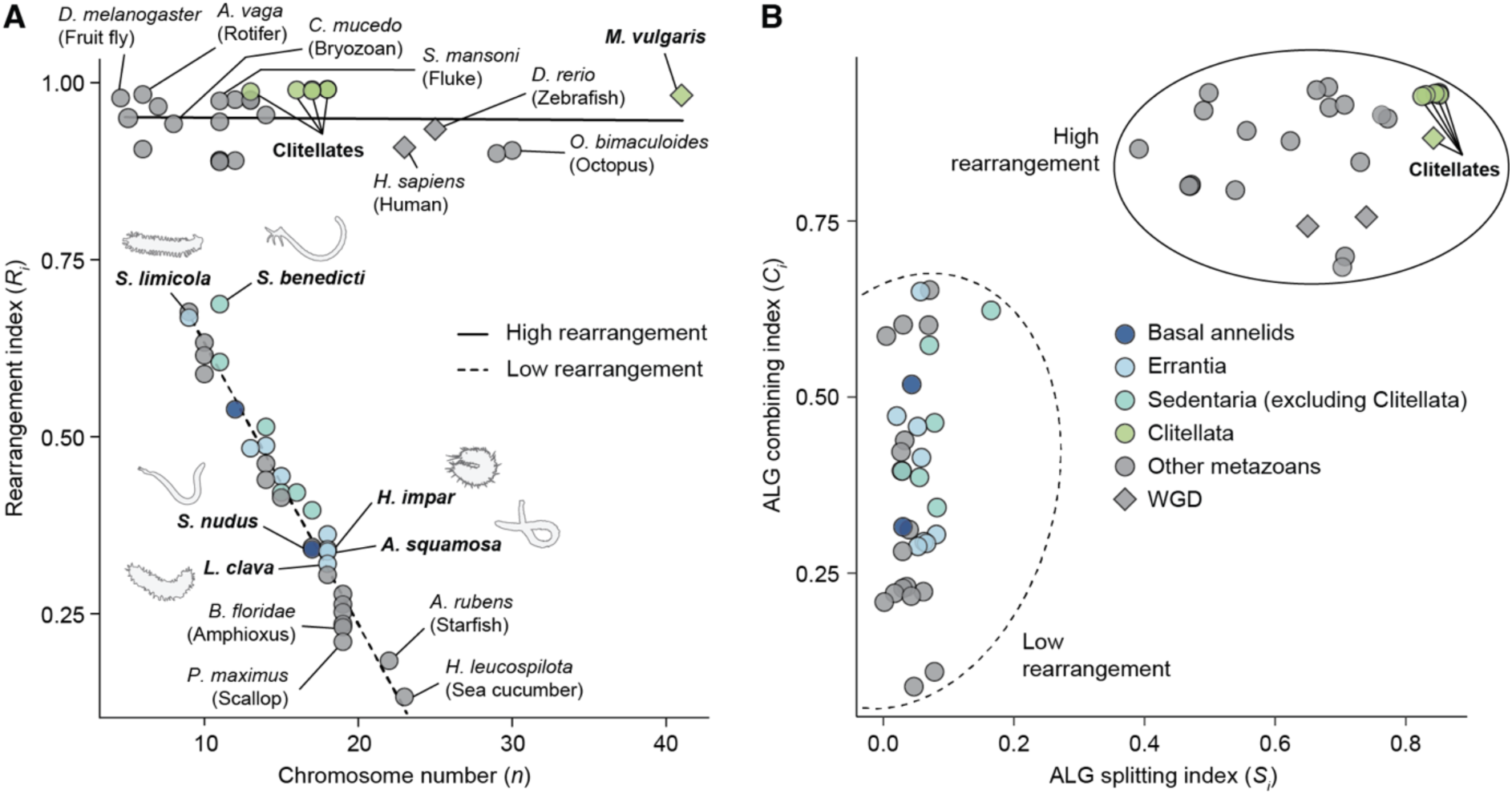
Clitellate annelids have unique levels of interchromosomal rearrangements among bilaterians. (**A**) Rearrangement index versus chromosome number in bilaterian genomes. Rearrangement index measures bilaterian ancestral linkage group fusion and splitting: higher indices reflect more rearrangement, and lower indices show less rearranged genomes. Lineages with whole genome duplications (WGD) are marked with a diamond. The relationships between rearrangement index and chromosome number were examined using linear models for high rearrangement (solid line) and low rearrangement (dashed line) groups. The estimated model for low rearrangement species *y = -0.040x + 1.032* has a statistically significant slope (SE = 0.002, *t* = -25.500, *P* = 7.378 x 10^-23^), suggesting that rearrangement index decreases linearly with chromosome number. The *R^2^* value of 0.956 suggests that approximately 96% of the variation in rearrangement index in these species is explained by chromosome number. The estimated model for high rearrangement species y = -0.0001x + 0.952 has a non-significant slope (SE = 0.001, *t* = -0.142, *P* = 0.888), suggesting that chromosome number does not predict rearrangement index in these species. The *R^2^* value of -0.039 suggests that no variation in the rearrangement index is predicted by chromosome number. (**B**) ALG combining index versus ALG splitting index in bilaterian genomes. The ALG splitting index (*S_i_*) = 1 - *S* (ALG splitting parameter). The ALG combining index (*C_i_*) = 1 - *C* (ALG combining parameter). See Materials and Methods for full description of the calculation of these parameters. Higher index values indicate increased levels of rearrangement.

Interestingly, we found that bilaterian genomes fell into one of two groups, a high rearrangement group and a low rearrangement group, which are distinctly separate on the plot (fig. 5A; supplementary fig. S15). We used linear models to examine the relationship between rearrangement index and chromosome number for each of the two groups (supplementary fig. S15). This showed that, in the low rearrangement group, over 95% of the variation of rearrangement index was explained by chromosome number (*P* = 7.378 x 10^-23^, *R^2^* = 0.956). Indeed, rearrangement index increases linearly as chromosome number decreases from the bilaterian ancestral state of 24 gene linkage groups. This suggests that the index largely reflects chromosome fusion. Non-clitellate annelids conform to this pattern: for instance, *L. clava* (*n* = 18) has the lowest rearrangement index and *Streblospio benedicti* (*n* = 11) the highest. In contrast, the chromosome number is not predictive of rearrangement index in the high rearrangement group (*P* = 0.888, *R^2^* = -0.039). This suggests that there are two separate groups of bilaterians in which different patterns of interchromosomal rearrangements are observed. Clitellate annelids are part of the high rearrangement group, while non-clitellate annelids fall into the low rearrangement group.

To better understand the factors distinguishing these two groups, we separated out the bilaterian ALG combining and ALG splitting parameters (fig. 5B; supplementary fig. S16). Our analysis reveals that both ALG combining and ALG splitting are higher in the high rearrangement group but that there is a large gap, particularly in the ALG splitting index. Low rearrangement group species have minimal ALG splitting, while the high rearrangement group has substantial levels of splitting. Based on the annelid genomes, we can infer that the difference between these two groups is the absence of chromosome fission in the low rearrangement species. To further understand the underlying causes of this distinction, rather than averaging the index across all ALGs, we performed a principal component analysis (PCA) using the individual index scores for each ALG in each species (supplementary fig. S17A, supplementary table S8). The high rearrangement and low rearrangement groups were well separated in the PCA. The dimension distinguishing the two groups (Dim-1) accounted for over 75% of the variance in the dataset, and the top 20 variables contributing to this dimension were all ALG splitting indices (supplementary fig. S17B). This confirms that the major factor separating the high and low rearrangement groups is the extent of ALG splitting. In turn, this suggests that there are two groups of bilaterian genomes, those in which ALG fission is evolutionarily permitted and frequent and those in which it is highly restricted.

### Macrosynteny as a tool for taxonomy within a phylum

The past year has seen macrosynteny emerge as a novel tool for delineating phylogenetic relationships (Parey et al. 2023; Schultz et al. 2023; Lewin et al. 2024; Steenwyk and King 2024). In particular, chromosome fusion-with-mixing events have high potential as phylogenetically informative rare genomic changes (i.e. molecular synapomorphies) because they are irreversible (Rokas and Holland 2000; Schultz et al. 2023; Steenwyk and King 2024). We used our dataset to test the power of ALG-based macrosynteny as a taxonomic tool by asking whether it can be used to reliably identify characteristics for defining monophyletic groups of annelids. Within the dataset of 16 non-clitellate species, we found four clades with lineage-defining interchromosomal rearrangements. First, all annelids except *Owenia fusiformis* have a C2⊗(J2⊗L) fusion; second, within Errantia, the suborder Aphroditiformia (scale worms) is defined by a C1⊗partial_(H⊗Q) fusion; third, members of the family Polynoidae share a further A1⊗E fusion; and fourth, the Sedentaria family Siboglinidae (giant tube worms) share four fusions (A2⊗M, B2⊗J1, B3⊗(O1⊗R), D⊗P) (fig. 6A). We note that several of these changes occur in a stepwise fashion, leading to progressively more derived genomes. For instance, all sampled annelids except *O. fusiformis* have C2⊗(J2⊗L); within this group, Aphroditiformia annelids have C1⊗partial_(H⊗Q); then within Aphroditiformia the Polynoidae have A1⊗E; and within Polynoidae there are species-specific changes (e.g. I⊗N, M⊗(K⊗O2) and partial_(H⊗Q)⊗(C2⊗(J2⊗L) in *Alentia gelatinosa*).

**Fig. 6.**
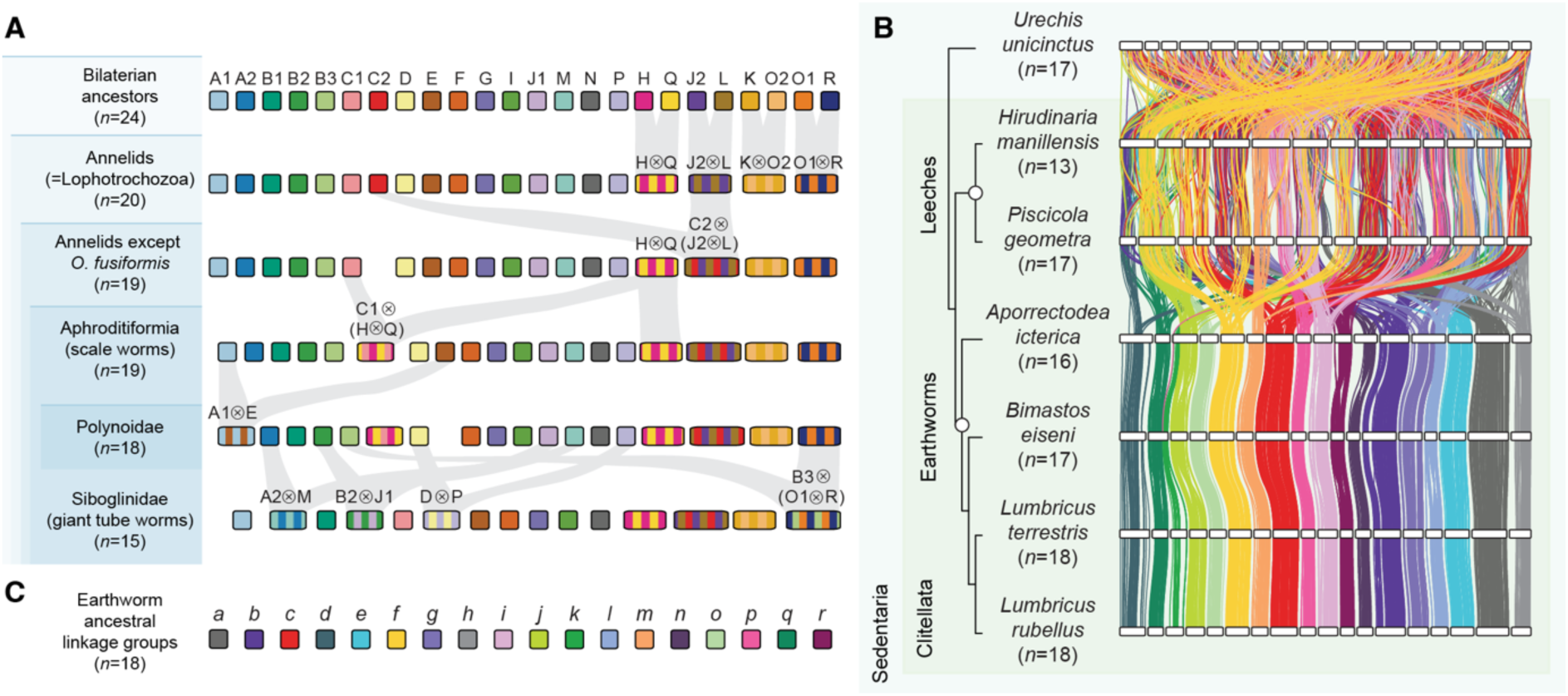
Conservation of interchromosomal rearrangements defines distinct taxonomic groups. (**A**) Stepwise evolution of genome reorganization events in Annelida. Boxes represent linkage groups or chromosomes colored by their bilaterian ancestral state. Fusion-with-mixing events are represented by striped chromosomes. Interchromosomal rearrangements that are diagnostic for specific lineages are highlighted in gray ribbons. Shaded areas on the left between taxa represent progressively smaller subsets of species. For instance, Polynoidae is a subset of Aphroditiformia. (**B**) Ideogram plots of clitellate genomes colored by earthworm ancestral linkage groups. Earthworm ALGs are defined by genes’ position in *Lumbricus rubellus.* (**C**) Color code for earthworm ALGs. These are highly conserved across the four earthworm species in this dataset.

These clade-defining rearrangements reveal the potential of ALG-based macrosynteny for within-phylum systematics. For instance, the absence of the C2⊗(J2⊗L) fusion in *O. fusiformis* unequivocally confirms the Oweniidae as a basal annelid lineage and Sipuncula as an annelid despite its lack of segmentation and appendages, more closely related to Errantia and Sedentaria than to *O. fusiformis*. This data demonstrates the ability of interchromosomal rearrangements to be used as taxonomically-informative characters.

To facilitate macrosynteny comparisons in the completely rearranged clitellate genomes, we assigned newly formed but subsequently conserved gene groups to ALGs. Determination of earthworm (fig. 6B and C; supplementary fig. S18) and leech (supplementary fig. S19) ALGs reveals strong conservation of macrosynteny within each group but also partial conservation of ALGs shared across both taxa. Therefore, leeches and earthworms can be defined by their own distinct ALGs, and the two groups share a recent common ancestor in which genome rearrangement had already begun. Overall, in this dataset alone, at least seven monophyletic groups (non-basal annelids, Aphroditiformia, Polynoidae, Siboglinidae, Clitellata, leeches, and earthworms) can be defined by specific interchromosomal rearrangements that may be used for taxonomic classification (supplementary table S9).

## Discussion

Our study reveals two distinct groups of annelids: non-clitellates, in which genome organization is characterized by broad conservation of the ancestral bilaterian genome architecture followed by limited chromosome fusions; and clitellates, in which the genome has been shuffled by fusion and fission to such an extent that bilaterian ALGs have been completely lost. Indeed, this is a microcosm of the situation within bilaterians as a whole. By quantifying the extent to which genomes are rearranged, we found that bilaterians, like annelids, fall into one of two categories: low rearrangement, typically with some ALG fusion but low levels of fission, and high rearrangement, in which both fusion and fission are common. This supports the hypothesis that there is an evolutionary constraint on genome structure (present in non-clitellates) which can be flicked off like a switch (clitellates), causing rapid, extensive genome rearrangement and the complete disintegration of previously conserved ALGs.

A key open question is what selective pressures control the switch that flicks from the conservative evolution of ALGs to their complete atomization in specific clades. Given that clitellates possess the highest rearrangement index of all species in our dataset, they may be an optimal group with which to answer this question. Previous studies have noted that the loss of bilaterian genome structure in clitellates coincides with the transition from marine to freshwater and terrestrial environments (Rousset et al. 2008; Erséus et al. 2020; Sun et al. 2021). Indeed, this process was associated with a high degree of morphological evolution in clitellates, including the presence of the cocoon-producing clitellum and a highly derived mechanism of development (Kuo 2017). Genome rearrangement may be selectively favored during adaptation to a radically different environment because it facilitates changes to regulatory landscapes and therefore novel gene expression patterns. Consistent with this, Hox genes, the expression of which is (a) critical for early development and (b) highly dependent on genome organization (Duboule 2007; Rekaik and Duboule 2024), are extensively rearranged in the genomes of the earthworms *Eisenia fetida* and *Perionyx excavatus* and the leech *H. robusta* (Cho et al. 2012; Simakov et al. 2013; Zwarycz et al. 2015; Barucca et al. 2016). Importantly, there are data suggesting that Hox expression is divergent in leeches compared to other annelids (Kourakis and Martindale 2001; Gąsiorowski et al. 2023), but a more comprehensive analysis is needed to confirm this. Overall, dramatic genome rearrangement in clitellates correlates with the evolution of a new ecological niche, alongside divergent genomic location and altered expression of key developmental genes.

Our phylum-level dataset is of sufficient depth to start to identify trends in ALG evolution. First, it reveals that species within a phylum can have completely divergent genome structures, with bilaterian ALGs preserved in some and completely lost in others. Second, it shows that interchromosomal rearrangements can occur both in a gradual stepwise fashion (non-clitellates) and as rapid, sweeping changes (clitellates). Third, ALG fusion is almost always followed by ALG mixing within the chromosome, and fusion-without-mixing is very rare. Fourth, fusion of ALGs is much more common than fission, suggesting strong selective pressures to maintain genes together on the same chromosome. This is supported by data from Lepidoptera (butterflies and moths), which also revealed fission to be much less common than fusion (Wright et al. 2024).

While methods for phylogeny reconstruction using microsynteny (small-scale conservation of gene order) are becoming increasingly sophisticated (Drillon et al. 2020; Zhao et al. 2021), the use of ALG-based macrosynteny for phylogenetic inferences is in its infancy. In general, though not without exception (Li et al. 2022), macrosynteny appears to decay slower than microsynteny (Simakov et al. 2022), meaning that it may have a unique utility for delineating relationships between distantly related groups. Recent works placing ctenophores as the basal metazoan lineage (Schultz et al. 2023), suggesting that bryozoans are closely related to brachiopods (Lewin et al. 2024) and resolving branching order in teleost fishes (Parey et al. 2023), highlight its significant potential. Within this dataset of 23 annelids, we describe unique chromosome rearrangements that can be used as rare genomic changes to define seven different taxonomic groups at levels varying from class to family. For example, genome structure definitively supports Sipuncula (in the past considered a separate phylum) as an annelid and Oweniidae as a basal annelid lineage due to the presence of the C2⊗(J2⊗L) fusion in the former and absence in the latter, confirming data from sequence-based phylogenetics (Weigert et al. 2014; Struck et al. 2015; Zhong et al. 2022; Zheng et al. 2023). Importantly, the observed stepwise manner of ALG rearrangements suggests that changes to genome structure as clade-defining characters need not be restricted to a specific taxonomic level but can be applied at any level from metazoan-wide to genus and species. At present, the sampling depth is likely to limit the utility of this to a few, specific cases, but the accelerating accumulation of chromosome-level assemblies makes it inevitable that in the coming years, many groups will have sufficiently dense sampling for robust genome structure-based taxonomic definitions.

One strength of this framework is its potential for disentangling the evolutionary relationships between fast-evolving lineages. Rapidly-evolving genomes like those of clitellates, while troublesome for sequence-based phylogenetics due to artifacts like long branch attraction (Felsenstein 1978; Bergsten 2005), may be ideal for genome-structure-based taxonomy due to the rapid accumulation of genome rearrangements and the improbability that highly complex rearrangements could be convergently evolved. Therefore, genome structure-based taxonomy may be particularly helpful for elucidating the positions of traditionally problematic lineages.

## Materials and Methods

### Assembly acquisition and gene prediction

This study aimed to characterize interchromosomal rearrangements within the phylum Annelida. All available chromosome-level assemblies of annelid species (*n* = 24) were obtained from the National Center for Biotechnology Information (NCBI) using NCBI Datasets on February 1st, 2024. Of the 24 genomes, 16 were produced by the Darwin Tree of Life (DToL) sequencing project (The Darwin Tree of Life Project Consortium et al. 2022). The genome assemblies from the DToL project have been made publicly available to the community for further analysis. Those with an accompanying publication are *Acholoe squamosa* (Adkins et al. 2023), *Alitta virens* (Fletcher et al. 2023), *Lepidonotus clava* (Darbyshire et al. 2022), *Piscicola geometra* (Doe et al. 2023), and *Sthenelais limicola* (Darbyshire et al. 2023). Genomes from other sources with accompanying publications are: *Branchipolynoe longqiensis* (He et al. 2023), *Hirudinaria manillensis* (Liu et al. 2023), *Metaphire vulgaris* (Jin et al. 2020), *Owenia fusiformis*, (Martín-Zamora et al. 2023), *Paraescarpia echinospica (Sun et al. 2021)*, *Streblospio benedicti* (Zakas et al. 2022), *Sipunculus nudus* (Zheng et al. 2023), and *Urechis unicinctus* (Cheng et al. 2024).

The genome for *Branchellion lobata* was excluded from the main analyses because it has a low BUSCO completeness score (72.9% with the metazoan_obd10 database) (Simão et al. 2015) but is included as a supplementary figure (supplementary fig. S20). This left a dataset of 23 species. One species, *O. fusiformis*, had available GenBank gene annotations. Gene prediction for the remaining 22 species was performed using RepeatModeler2 (v2.0.4) (Flynn et al. 2020), RepeatMasker (v4.1.5) (Smit et al. 2015), and the BRAKER3 pipeline (v3.0.3) (Stanke et al. 2006; Stanke et al. 2008; Li et al. 2009; Barnett et al. 2011; Lomsadze et al. 2014; Buchfink et al. 2015; Hoff et al. 2016; Hoff et al. 2019; Brůna et al. 2021) as reported previously (Lewin et al. 2024). For species with available RNA-seq data (supplementary table S10), reads were trimmed with fastp (v0.23.4) (Chen et al. 2018) and mapped with STAR (v2.7.10b) (Dobin et al. 2013) before BRAKER3 was run in RNA-seq mode. For species with no RNA-seq data, BRAKER3 was run in protein mode using the supplied Metazoa.fa protein file. Gene prediction quality was assessed using BUSCO (v5.4.7) (Simão et al. 2015).

### Phylogenetic analysis

Single-copy orthologs were identified with OrthoFinder (v2.5.4) (Emms and Kelly 2019). The tree splitting and pruning algorithm of OrthoSNAP (v0.0.1) (Steenwyk et al. 2022) was then used to recover additional single-copy orthologs from gene family trees. Sequences of each ortholog were aligned with MAFFT (v7.520) (Katoh et al. 2002; Katoh and Standley 2013), trimmed with ClipKIT (v1.4.1) (Steenwyk et al. 2020) and concatenated with PhyKIT (v1.11.7) (Steenwyk et al. 2021), before maximum likelihood phylogeny inference with IQ-TREE (v2.2.2.3) (Minh et al. 2020). ModelFinder (Kalyaanamoorthy et al. 2017) was used for automatic substitution model selection, and UFBoot2 (Hoang et al. 2018) was used to perform 1,000 ultra-fast bootstrap replicates.

### Macrosynteny analysis

SyntenyFinder (Lewin et al. 2024) was used to implement OrthoFinder (Emms and Kelly 2019) and RIdeogram (v0.2.2) (Hao et al. 2020) and produce Oxford dot plots. Bilaterian ALGs were determined by orthology (Simakov et al. 2022). Unless stated otherwise, links between chromosomes with fewer than 10 shared orthologs are trimmed from ribbon plots for clarity; all genes are shown in Oxford dot plots.

### Whole genome duplication inference

Several complementary methods were used to test for WGDs: 1) structural information from ideogram plots and Oxford dot plots using single-copy orthologs as above; 2) structural information from Oxford dot plots using multi-copy orthologs, permitting up to five paralogs in one species; 3) *K_S_* plots showing distributions of synonymous substitutions per synonymous site, produced using ‘wgd dmd’ and ‘wgd ksd’ in wgd (v2.0.26) (Zwaenepoel and Van de Peer 2019; Chen and Zwaenepoel 2023); 4) identification of blocks of genes with conserved microsynteny within annelid genomes using ‘wgd syn’ (Zwaenepoel and Van de Peer 2019; Chen and Zwaenepoel 2023).

### Rearrangement index

A ‘rearrangement index’ (*R_i_*) was developed to quantify the extent to which bilaterian ALG rearrangement has occurred in bilaterian genomes. For each ALG, the rearrangement index is calculated as:

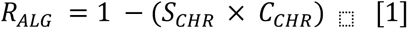

where *R_ALG_* denotes the rearrangement index for a given ALG; *S_CHR_* (ALG splitting parameter) represents the highest proportion of genes from this ALG on a single chromosome and *C_CHR_* (ALG combining parameter) is the proportion of genes on that chromosome that belong to that particular ALG. By incorporating these parameters, the index accounts for both ALG splitting and ALG combining.

Subsequently, the *R_i_* for each genome is given by the equation:

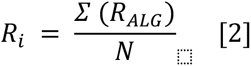

where *R_i_* denotes the rearrangement index for the genome; *R_ALG_* the is rearrangement index for each ALG; and *N* is the total number of ALGs. The higher the index, the higher the level of interchromosomal rearrangements.

### RNA-sequencing data analysis

Raw RNA-seq data (supplementary table S11) from *M. vulgaris* tissues were downloaded from NCBI SRA using SRA toolkit (Leinonen et al. 2011) and GNU parallel (v20230322) (Tange 2021). Gene expression was quantified with the pseudo-aligner salmon (v1.10.2) (Patro et al. 2017).

### Statistical analysis

Statistical analysis was performed using R (v4.3.0) (R Core Team 2023). Spearman’s rank correlation was used to test whether the number of fusions of each ALG correlates with the average length of chromosomes on which the ALG is hosted, as described previously (Wright et al. 2024). Chromosome length was measured as a proportion of the total genome length. Chi-squared tests were used to test for differences in the number of fusions per ALG and the number of fusions per species. Linear models were used to examine the relationship between rearrangement index and chromosome number. *P* < 0.05 was considered to be statistically significant.

## Supporting information

ESM1 - Supplementary Figures

ESM2 - Supplementary Tables

## Acknowledgments

This work was partially supported by a Royal Society Newton International Fellowship (NIF\R1\201315) to Y.-J.L. and an Academia Sinica Career Development Award (AS-CDA- 112-L06) to Y.-J.L. We thank the members of the Symbiosis Genomics & Evolution Lab for their assistance and support and Stephan Q. Schneider for thoughtful discussions and advice.

## Author Contributions

T.D.L. and Y.-J.L. conceived the project. T.D.L. annotated the genomes; T.D.L., I.J.-Y.L., and Y.-J.L. developed the bioinformatics pipeline; T.D.L. analyzed the data; T.D.L and Y.-J.L. wrote the manuscript with input from I.J.-Y.L.

## Data Availability

All data are available in the main text or the supplementary materials.

