## Supplementary material for "Annelid comparative genomics and the evolution of massive lineage-specific genome rearrangement in bilaterians": ESM1 - Supplementary Figures

**This PDF file includes**

Supplementary Fig. S1 – S20

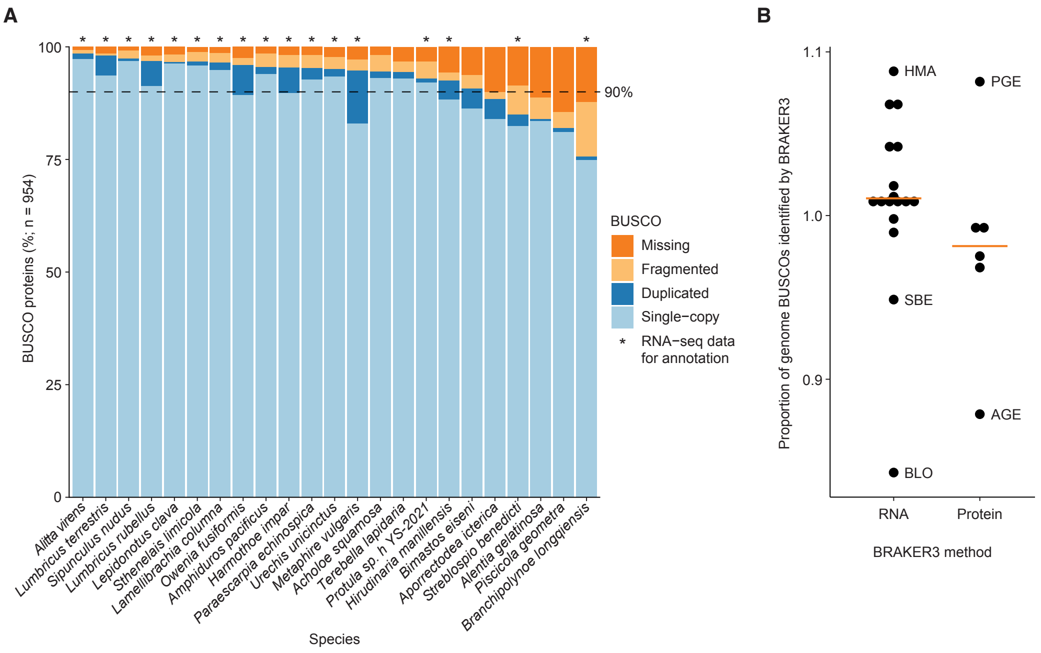


**Fig. S1. Gene prediction in annelid genomes.** (**A**) BUSCO metazoa *odb10* results for protein models in annelid genomes. Asterisks (*) mark species in which RNA-seq data was available to guide gene prediction. (**B**) Proportion of BUSCOs identified in the genome that appear in gene model predictions. Orange lines mark the mean. A score over 1 indicates that more BUSCOs were recovered from the gene model predictions than the raw genome sequence.

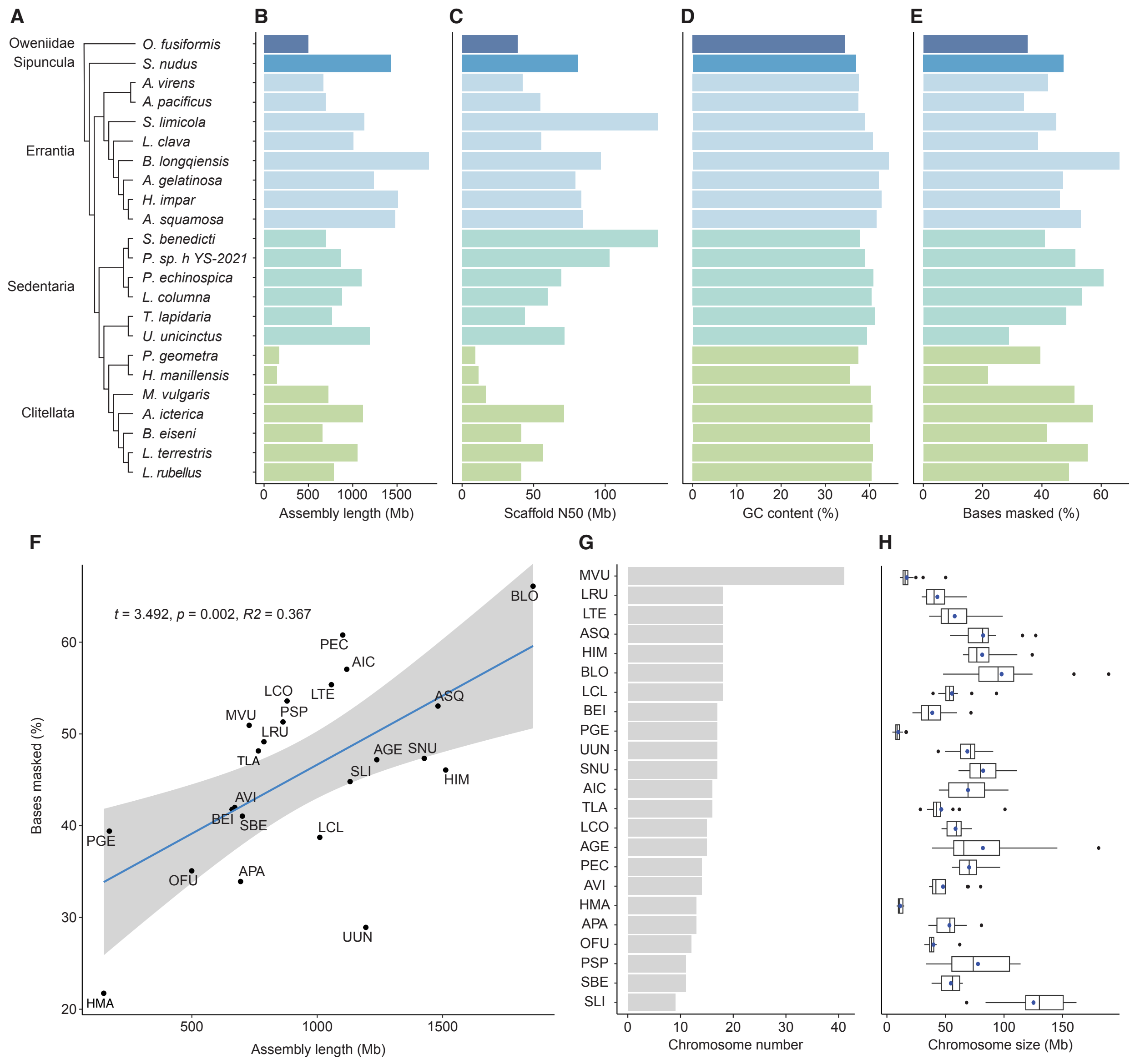


**Fig. S2. Assembly statistics for annelid genomes.** (**A**) Phylogeny of sampled annelids replicated from main text Fig. 1. (**B**) Assembly length. (**C**) Scaffold N50. (**D**) GC content. (**E**) Proportion of bases masked. (**F**) Proportion of bases masked increases with assembly length. (**G**) Chromosome number. (**H**) Chromosome size.

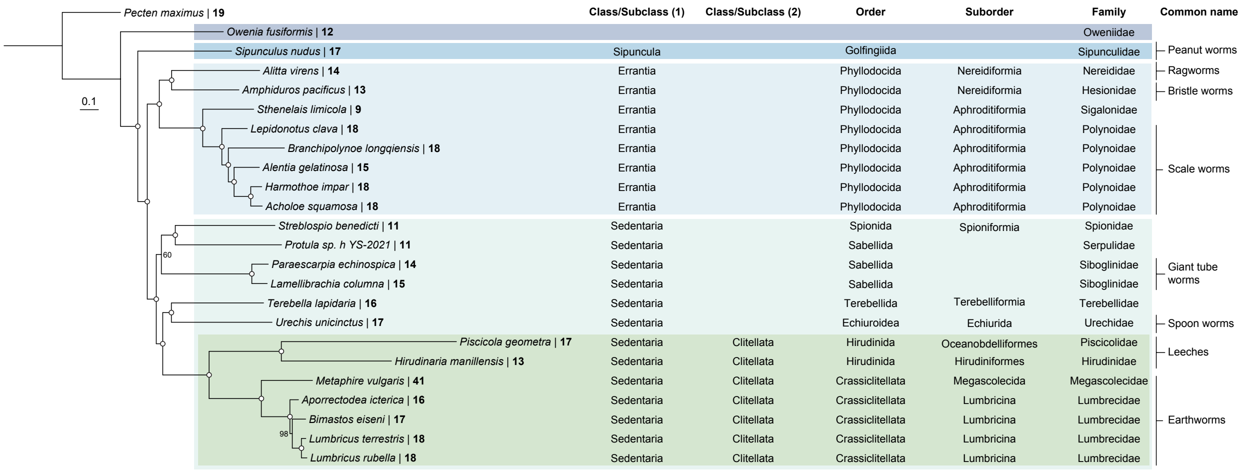


**Fig. S3. Phylogeny of annelids including taxonomic ranks.** Taxonomic classifications are drawn from NCBI Taxonomy and World Register of Marine Species (WoRMS).

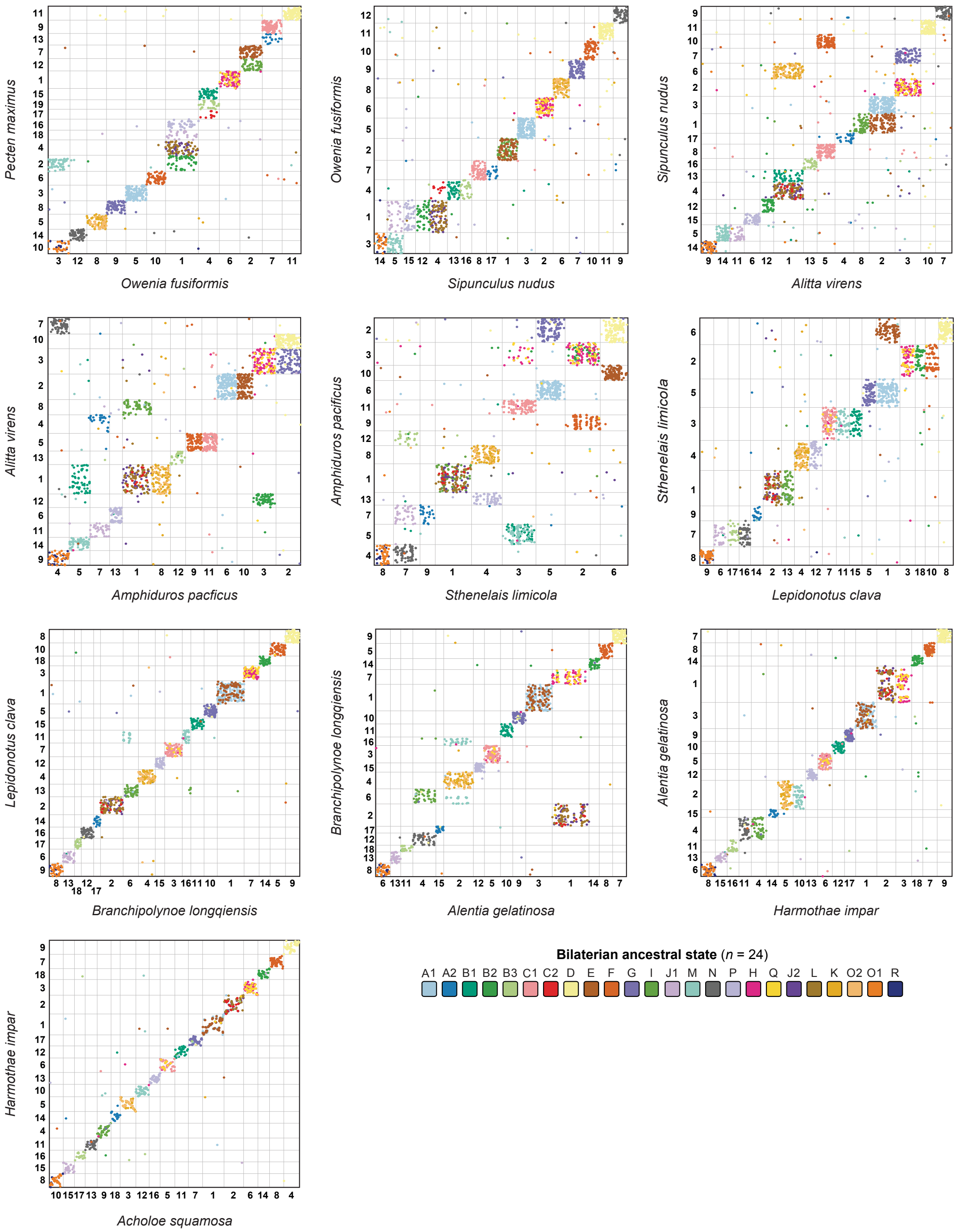


**Fig. S4. Oxford dot plots for Errantia annelids.** Each plot is a pairwise comparison between species, with each axis representing the full length of the species' genome. Each dot represents the location of an orthologous gene in the two species. Dots are colored by ALG. Integers represent chromosome numbers.

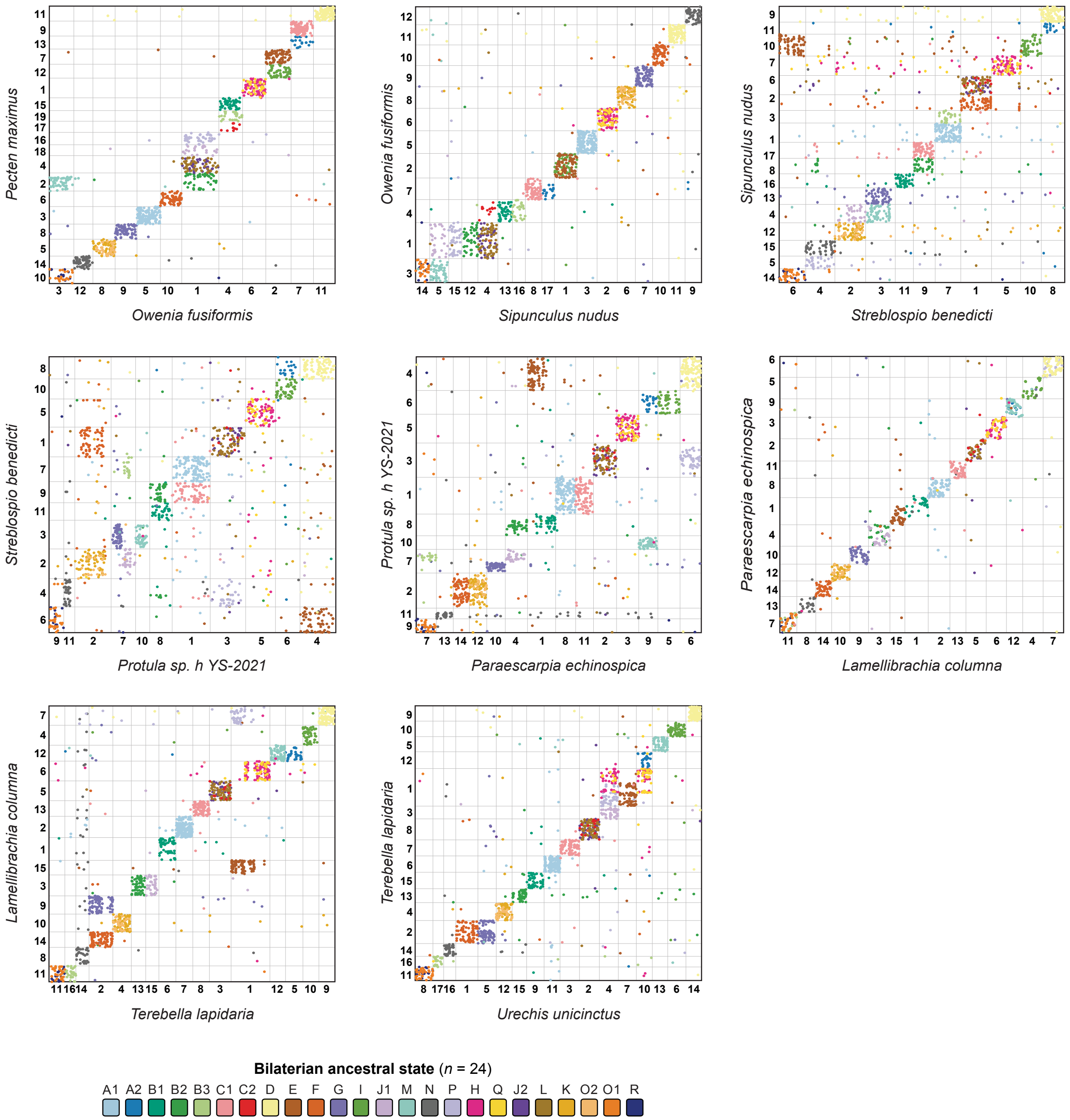


**Fig. S5. Oxford dot plots for Sedentaria annelids.** Each plot is a pairwise comparison between species, with each axis representing the full length of the species' genome. Each dot represents the location of an orthologous gene in the two species. Dots are colored by ALG. Integers represent chromosome numbers.

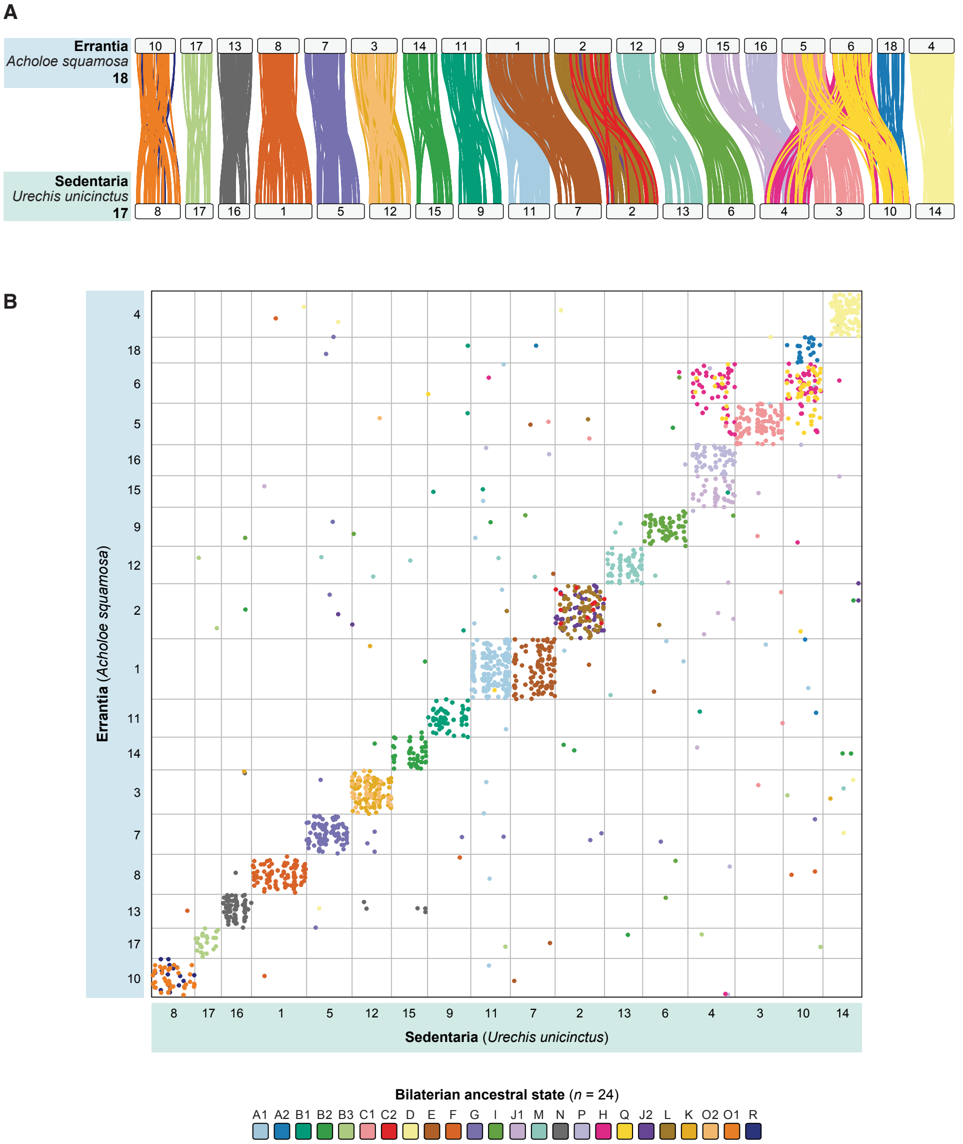


**Fig. S6. Genome structure comparison between Errantia and Sedentaria species.** (**A**) Ideogram plot for *Acholoe squamosa* (Errantia) *and Urechis unicinctus* (Sedentaria). Each line represents one pair of orthologues, colored by their ALG. (**B**) Oxford dot plot for *Acholoe squamosa* (Errantia) *and Urechis unicinctus* (Sedentaria). Plot is a pairwise comparison between species, with each axis representing the full length of the species' genome. Each dot represents the location of an orthologous gene in the two species. Dots are colored by ALG. Integers represent chromosome numbers.

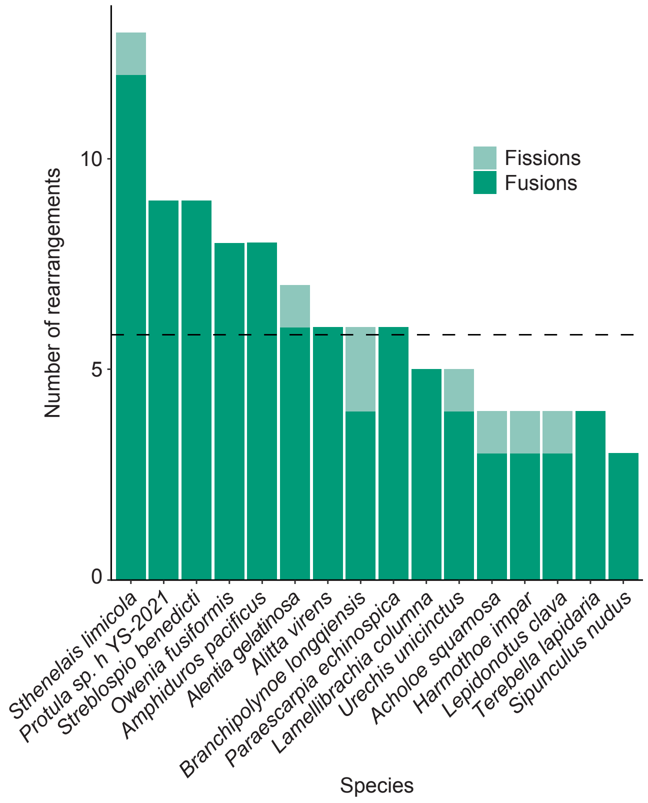


**Fig. S7. Number of chromosome rearrangements in annelid genomes.** A chi-squared test found no significant difference in the number of chromosome fusion across annelids ($P$ = 0.214, df = 15).

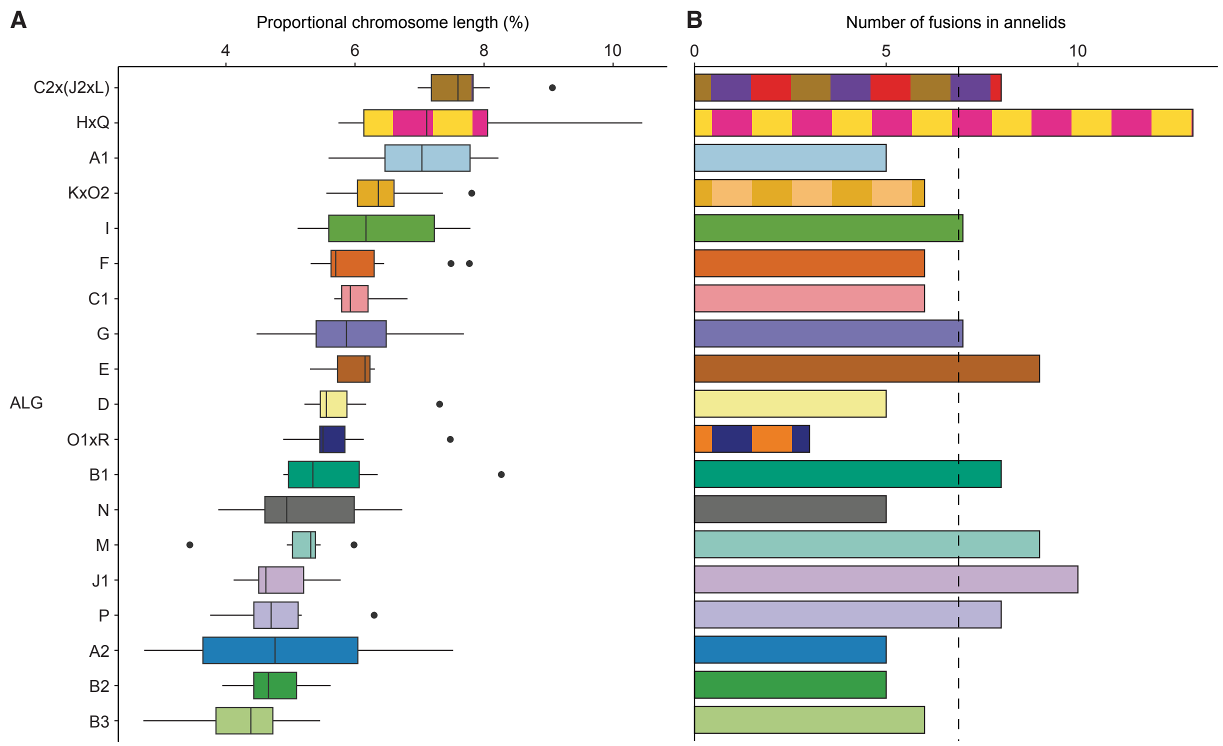


**Fig. S8. Rate of ALG fusion versus chromosome length.** (**A**) Length of chromosomes containing each ALG. Only chromosomes with no fusion or fission events are considered. (**B**) Number of fusions per ALG. Dotted line marks the mean. Spearman's rank correlation $\rho(19) = 0.125$, $P = 0.607$. A chi-squared test found no significant difference in the number of fusion events per ALG ($P = 0.736$, df = 15).

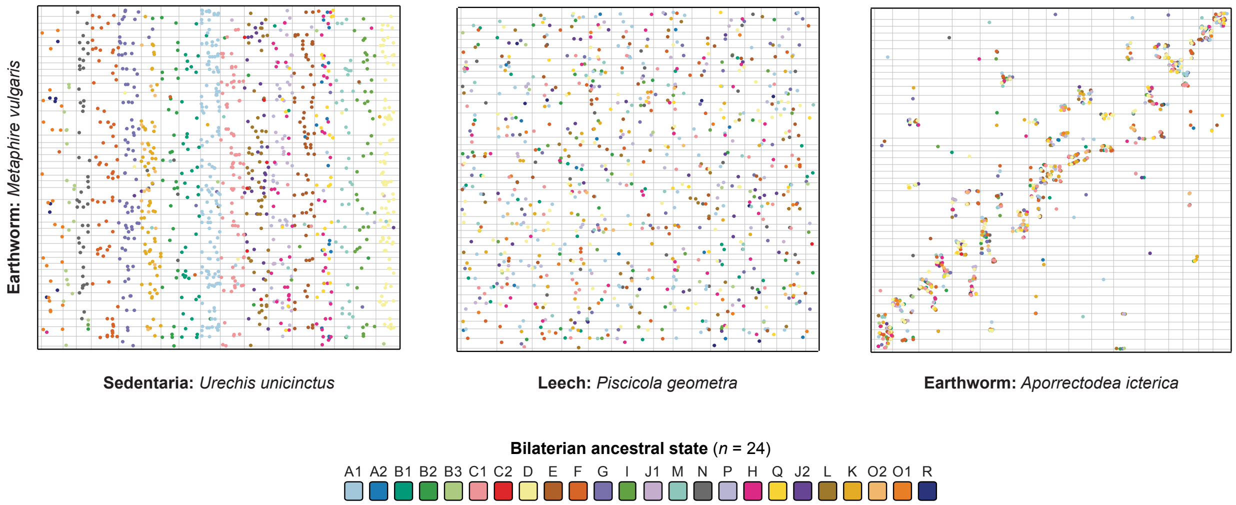


**Fig. S9. Oxford dot plots for the earthworm *Metaphire vulgaris*.** *M. vulgaris* has 41 chromosomes, at least 22 more than any other annelid sampled in this work. Its genome is completely shuffled compared to Sedentaria species such as *Urechis unicinctus*, and highly shuffled versus the leech *Piscicola geometra*. It shows partial conservation with the earthworm *Aporrectodea icterica*, but frequently with a 1:2 ratio of *A. icterica* genome sections to *M. vulgaris*. This is suggestive of whole genome duplication, which is explored further in main text Fig. 4 and Supplementary Fig. S10 – S13.

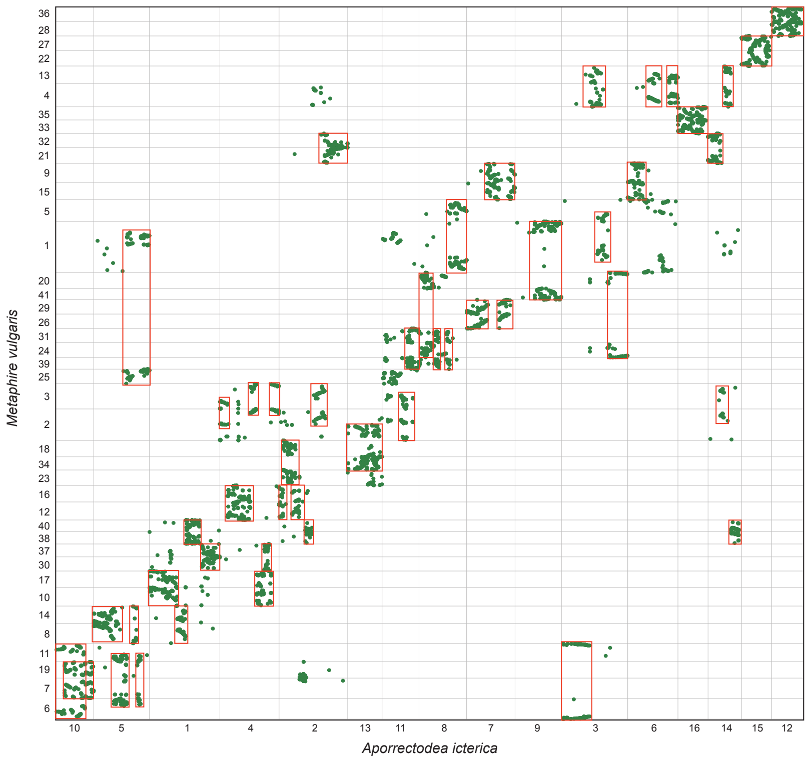


**Fig. S10. Oxford dot plot for orthologues between the earthworms *Metaphire vulgaris* and *Aporrectodea icterica.*** Plot is an enlarged version of main text Fig. 4B with chromosomes labelled and areas of 1:2 correspondence between *A. icterica* and *M. vulgaris* genomes highlighted with red boxes. This plot shows not only single-copy orthologues but also one:many orthologues with up to five copies in one species. The plot shows that many chromosome sections in *A. icterica* correspond to two chromosome sections in *M. vulgaris*, suggestive of whole genome duplication.

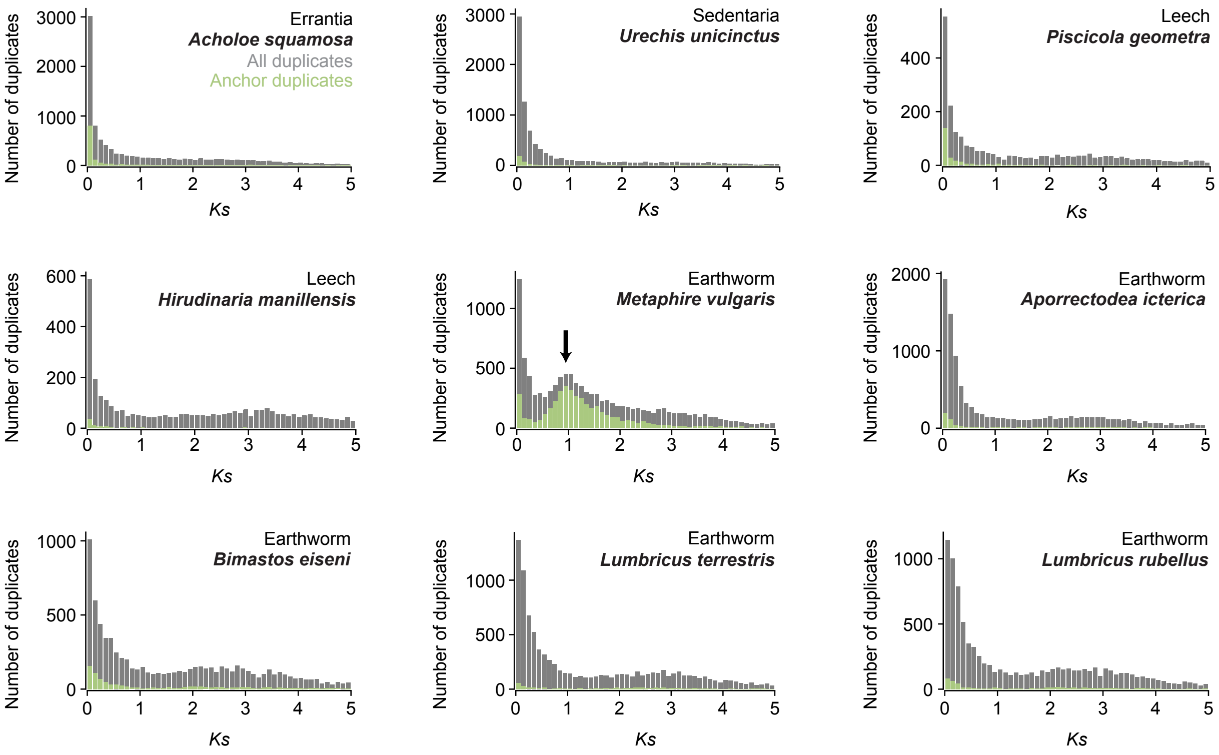


**Fig. S11. $K_s$ plots for annelid genomes.** Grey bars show histograms for all duplicated genes, while green bars show only anchor duplicates. Anchor duplicates are genes found in duplicated collinear blocks of genes in the genome. Genomes with no whole genome duplication are expected to show exponential decay of duplicate number as $K_s$ increases, as seen in all plots except that of *Metaphire vulgaris*. The *M. vulgaris* plot is interrupted by a normally distributed peak, indicative of whole genome duplication.

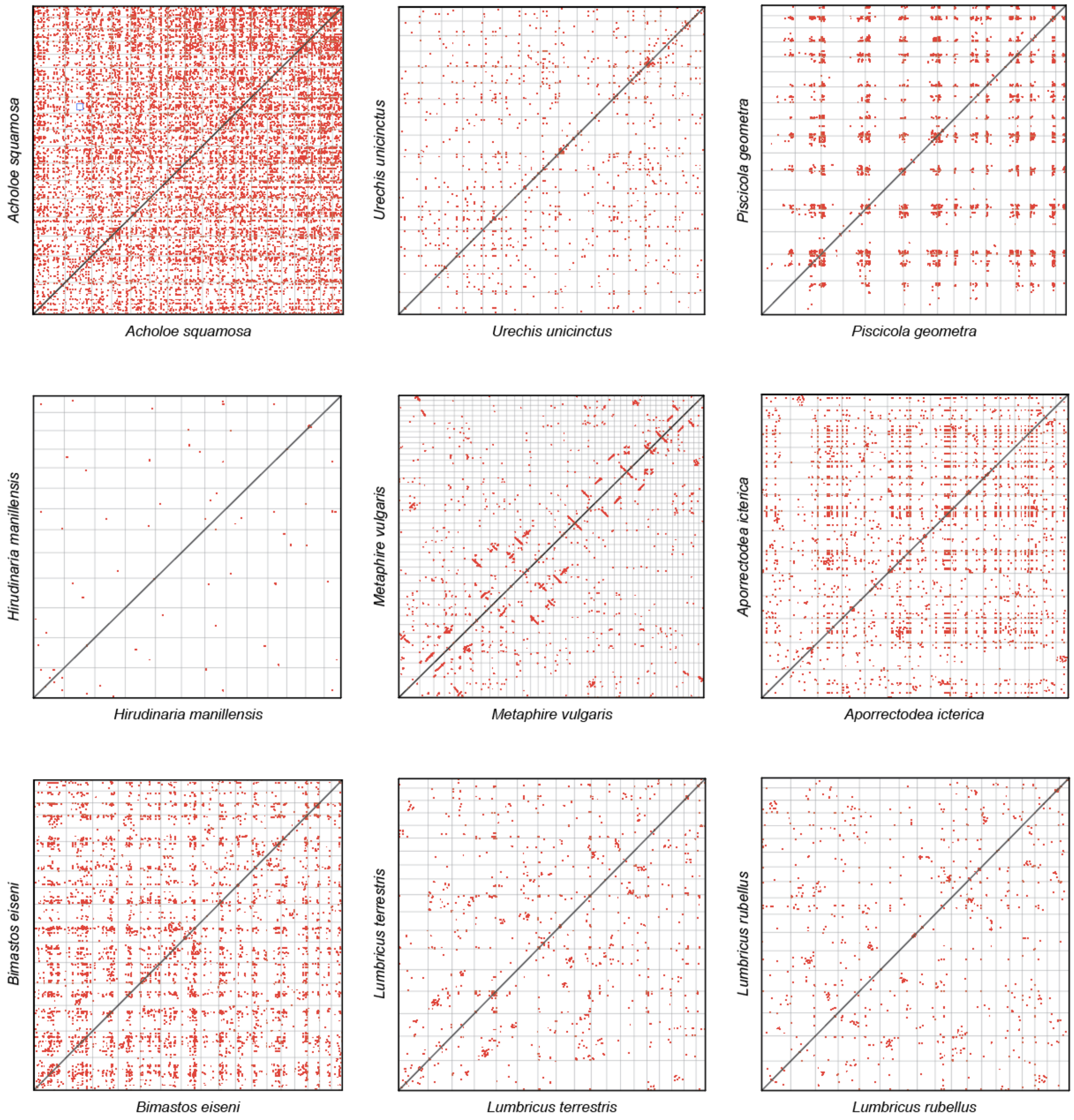


**Fig. S12. Intraspecific microsynteny conservation in annelid genomes.** Plots show instances of collinear blocks of homologous genes within a genome. The presence of many such blocks is suggestive of whole genome duplication. *Metaphire vulgaris* (center) is the only species with many such blocks (appearing as diagonal lines in the plot), suggesting that this species alone has undergone a recent whole genome duplication event. A full-size version of the *M. vulgaris* plot complete with chromosome labels is presented as Figure S13.

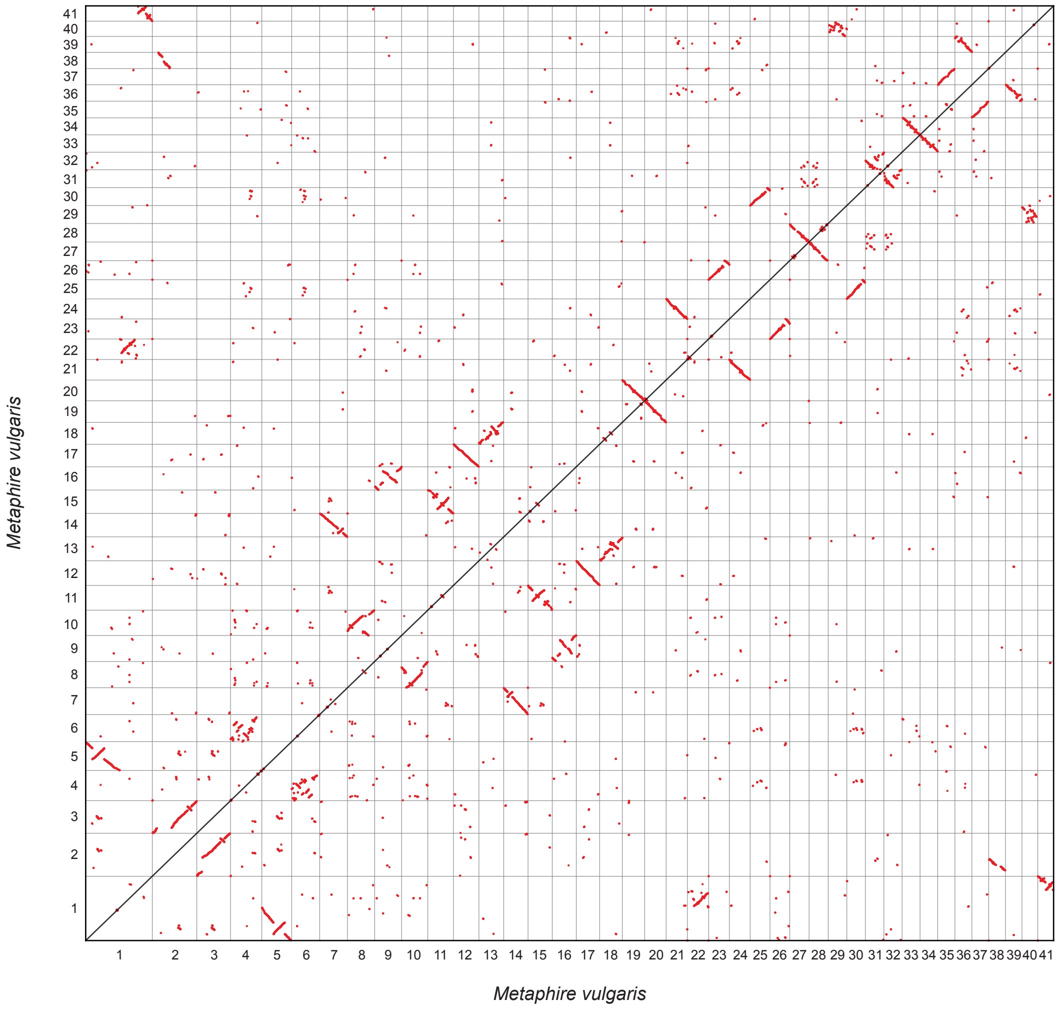


**Fig. S13. Intraspecific microsynteny conservation in *Metaphire vulgaris*.** Plot shows instances of collinear blocks of homologous genes within a genome: the presence of many such blocks is suggestive of whole genome duplication. *Metaphire vulgaris* is the only species with many such blocks (appearing as diagonal lines in the plot), suggesting that this species alone has undergone a recent whole genome duplication event.

**A. No rearrangement**

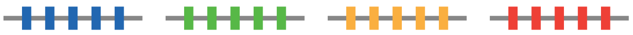

Rearrangement index = 0.00

| | Blue | Green | Yellow | Red | Mean |
|---|---|---|---|---|---|
| Splitting parameter (*S*) | 1 | 1 | 1 | 1 | 1 |
| Combining parameter (*C*) | 1 | 1 | 1 | 1 | 1 |
| Rearrangement index (1 - (*S* x *C*)) | 0 | 0 | 0 | 0 | 0 |

**B. Limited splitting**

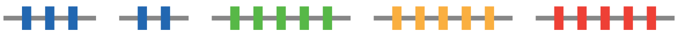

Rearrangement index = 0.10

| | Blue | Green | Yellow | Red | Mean |
|---|---|---|---|---|---|
| Splitting parameter (*S*) | 0.6 | 1 | 1 | 1 | 0.9 |
| Combining parameter (*C*) | 1 | 1 | 1 | 1 | 1 |
| Rearrangement index (1 - (*S* x *C*)) | 0.4 | 0 | 0 | 0 | 0.1 |

**C. Extensive splitting**

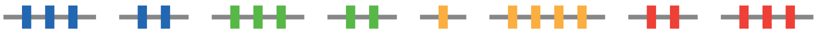

Rearrangement index = 0.35

| | Blue | Green | Yellow | Red | Mean |
|---|---|---|---|---|---|
| Splitting parameter (*S*) | 0.6 | 0.6 | 0.8 | 0.6 | 0.65 |
| Combining parameter (*C*) | 1 | 1 | 1 | 1 | 1 |
| Rearrangement index (1 - (*S* x *C*)) | 0.4 | 0.4 | 0.2 | 0.4 | 0.35 |

**D. Limited combining (similar to non-clitellate annelids)**

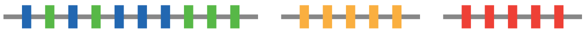

Rearrangement index = 0.25

| | Blue | Green | Yellow | Red | Mean |
|---|---|---|---|---|---|
| Splitting parameter (*S*) | 1 | 1 | 1 | 1 | 1 |
| Combining parameter (*C*) | 0.5 | 0.5 | 1 | 1 | 0.75 |
| Rearrangement index (1 - (*S* x *C*)) | 0.5 | 0.5 | 0 | 0 | 0.25 |

**E. Extensive combining**

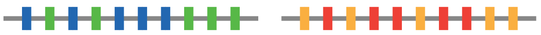

Rearrangement index = 0.50

| | Blue | Green | Yellow | Red | Mean |
|---|---|---|---|---|---|
| Splitting parameter (*S*) | 1 | 1 | 1 | 1 | 1 |
| Combining parameter (*C*) | 0.5 | 0.5 | 0.5 | 0.5 | 0.5 |
| Rearrangement index (1 - (*S* x *C*)) | 0.5 | 0.5 | 0.5 | 0.5 | 0.5 |

**F. Limited splitting and combining**

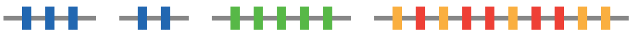

Rearrangement index = 0.33

| | Blue | Green | Yellow | Red | Mean |
|---|---|---|---|---|---|
| Splitting parameter (*S*) | 0.6 | 1 | 1 | 1 | 0.9 |
| Combining parameter (*C*) | 1 | 1 | 0.5 | 0.5 | 0.75 |
| Rearrangement index (1 - (*S* x *C*)) | 0.4 | 0 | 0.5 | 0.5 | 0.33 |

**G. Extensive splitting and combining (similar to clitellate annelids)**

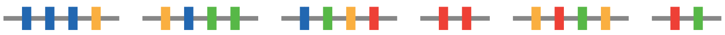

Rearrangement index = 0.69

| | Blue | Green | Yellow | Red | Mean |
|---|---|---|---|---|---|
| Splitting parameter (*S*) | 0.6 | 0.4 | 0.4 | 0.4 | 0.45 |
| Combining parameter (*C*) | 0.75 | 0.5 | 0.5 | 1 | 0.69 |
| Rearrangement index (1 - (*S* x *C*)) | 0.55 | 0.8 | 0.8 | 0.6 | 0.69 |

**Fig. S14. Explanation of the rearrangement index.** Parts (**A**) to (**F**) each show a highly simplified example of how the rearrangement index is calculated under different amounts of ALG splitting and ALG combining. Grey horizontal lines represent chromosomes and colored vertical lines represent genes. In this scenario there are four ALGs (blue, green, yellow, red) and each has 5 genes. This does not vary in any of the examples.

The rearrangement index is defined for each ALG as:

$$R_{ALG} = 1 - (S_{CHR} \times C_{CHR}) \quad [1]$$

where $R_{ALG}$ denotes the rearrangement index for a given ALG; $S_{CHR}$ (ALG splitting parameter) represents the highest proportion of genes from this ALG on a single chromosome and $C_{CHR}$ (ALG combining parameter) is the proportion of genes on that chromosome that belong to that particular ALG.

Then, the rearrangement index for each genome is defined as:

$$R_i = \frac{\Sigma(R_{ALG})}{N} \quad [2]$$

where $R_i$ denotes the rearrangement index for the genome; $R_{ALG}$ the is rearrangement index for each ALG; and $N$ is the total number of ALGs. Therefore, the rearrangement index for a given genome is the mean of the rearrangement indices for all ALGs in that genome.

A high rearrangement index indicates many interchromosomal rearrangements while a low index indicates few rearrangements.

For instance, the $R_{ALG}$ for example A (no rearrangements) blue ALG is calculated as follows: 5/5 blue genes are on one chromosome, so $S_{CHR}$ = 1; then, on this chromosome, 5/5 genes are blue genes so $C_{CHR}$ = 1. $R_{ALG}$ = 1 – (1 × 1) = 0. In example A, this is true for all four ALGs. So $R_i$ = (0 + 0 + 0 + 0) / 4 = 0.

In example B (ALG splitting by chromosome fission), the chromosome containing the blue ALG has fissioned into two parts. In this case, 3/5 blue genes are on one chromosome, so $S_{CHR}$ = 0.6; then, on this chromosome, 3/3 genes are blue genes (because there has been no fusion) so $C_{CHR}$ = 1. $R_{ALG}$ = 1 – (0.6 × 1) = 0.4. In example B, the other three ALGs have $R_{ALG}$ = 0, so $R_i$ = (0.4 + 0 + 0 + 0) / 4 = 0.1.

In example D (ALG combining by chromosome fusion), the chromosomes containing the blue and green ALGs have fused. For all ALGs $S_{CHR}$ = 1 because all genes are on the same chromosome. But, $C_{CHR}$ = 0.5 for the green and blue ALGs because only 5/10 of the genes on this new chromosome are from each one. Therefore, for green and blue ALGs, $R_{ALG}$ = 1 – (0.5 × 1) = 0.5. For red and yellow ALGs, $R_{ALG}$ = 1 – (1 × 1) = 0. Then, $R_i$ = (0.5 + 0.5 + 0 + 0) / 4 = 0.25.

Example G is the most complicated scenario, including extensive ALG splitting and combining similar to that observed in clitellate annelids. We will take the ALGs one by one. For the blue ALG, 3/5 genes are on one chromosome, so $S_{CHR}$ = 0.6. On this chromosome, 3/4 genes are blue, so $C_{CHR}$ = 0.75. $R_{ALG}$ = 1 – (0.6 × 0.75) = 0.55. For the green ALG, 2/5 genes are on one chromosome, so $S_{CHR}$ = 0.4. On this chromosome, 2/4 genes are green, so $C_{CHR}$ = 0.5. $R_{ALG}$ = 1 – (0.4 × 0.5) = 0.8. For the yellow ALG, 2/5 genes are on one chromosome, so $S_{CHR}$ = 0.4. On this chromosome, 2/4 genes are yellow, so $C_{CHR}$ = 0.5. $R_{ALG}$ = 1 – (0.4 × 0.5) = 0.8. Finally, for the red ALG, 2/5 genes are on one chromosome, so $S_{CHR}$ = 0.4. On this chromosome, 2/2 genes are red, so $C_{CHR}$ = 0.4. $R_{ALG}$ = 1 – (0.4 × 1) = 0.6. Then, $R_i$ = (0.55 + 0.8 + 0.8 + 0.6) / 4 = 0.69. This represents the highest rearrangement index in this simple set of scenarios.

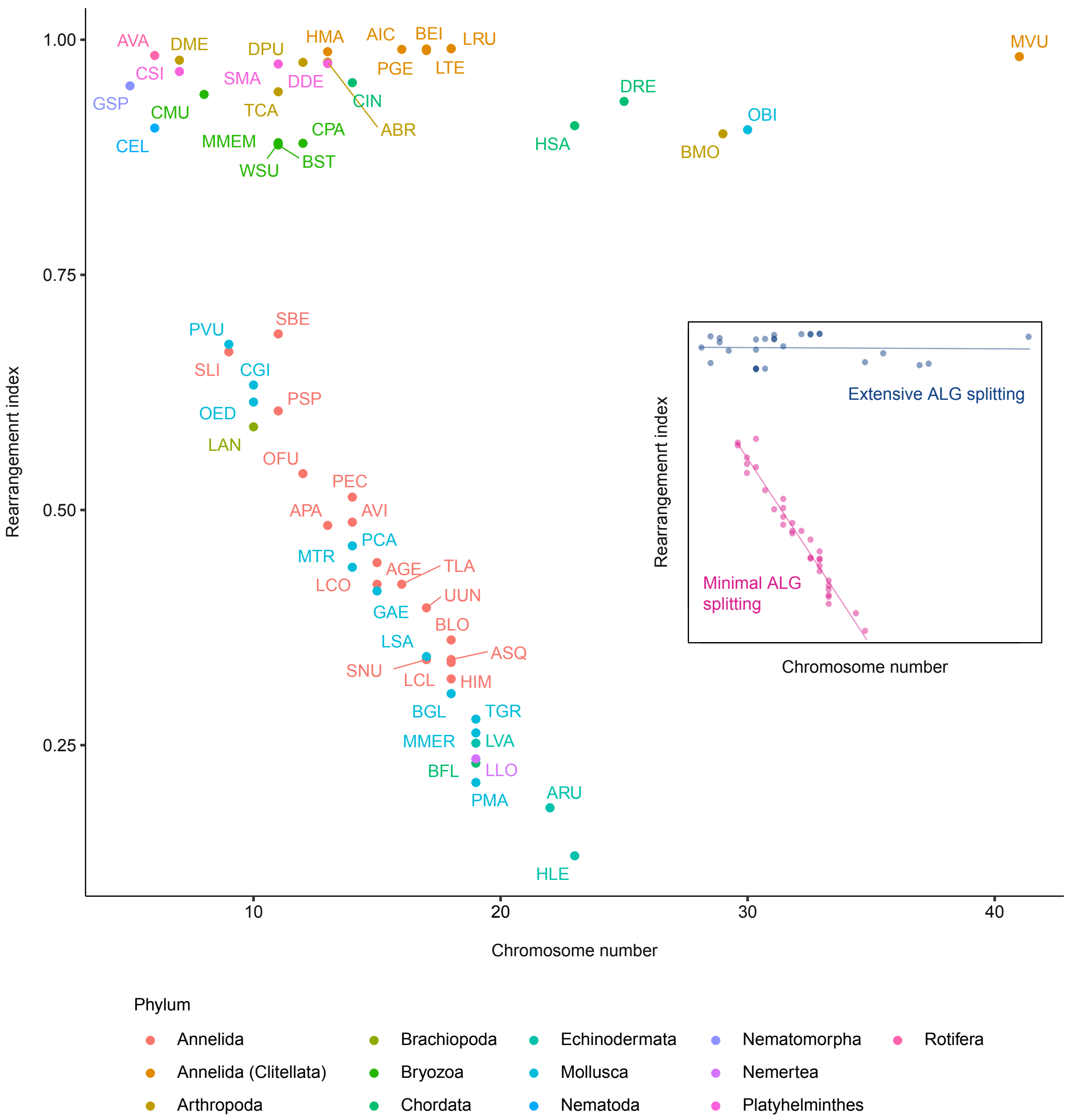


**Fig. S15. Bilaterians can be split into two groups based on their levels of interchromosomal rearrangements.** Plot is identical to that in main text Fig. 5A but with points colored by phylum. Rearrangement index is a measure of ALG combining (e.g. fusion) and ALG splitting (e.g. fission): higher rearrangement indices reflect more rearrangement and lower indices show less rearranged genomes. The plot reveals two distinct groups: those with a rearrangement index > 0.8 (including clitellates), and those with a rearrangement index < 0.7 (including non-clitellate annelids). Inset: the relationships between rearrangement index and chromosome number were examined using linear models for high rearrangement (blue) and low rearrangement (pink) groups. The estimated model for low rearrangement species $y = -0.040x + 1.032$ has a statistically significant slope (SE = 0.002, $t$ = -25.500, $P$ = 7.378 x $10^{-23}$), suggesting that rearrangement index decreases linearly with chromosome number. The $R^2$ value of 0.956 suggests that approximately 96% of the variation in rearrangement index in these species is explained by chromosome number. The estimated model for high rearrangement species y = -0.0001x + 0.952 has a non-significant slope (SE = 0.001, $t$ = -0.142, $P$ = 0.888), suggesting that chromosome number does not predict rearrangement index in these species. The $R^2$ value of -0.039 suggests that no variation in the rearrangement index is predicted by chromosome number. Species abbreviations are in table S7.

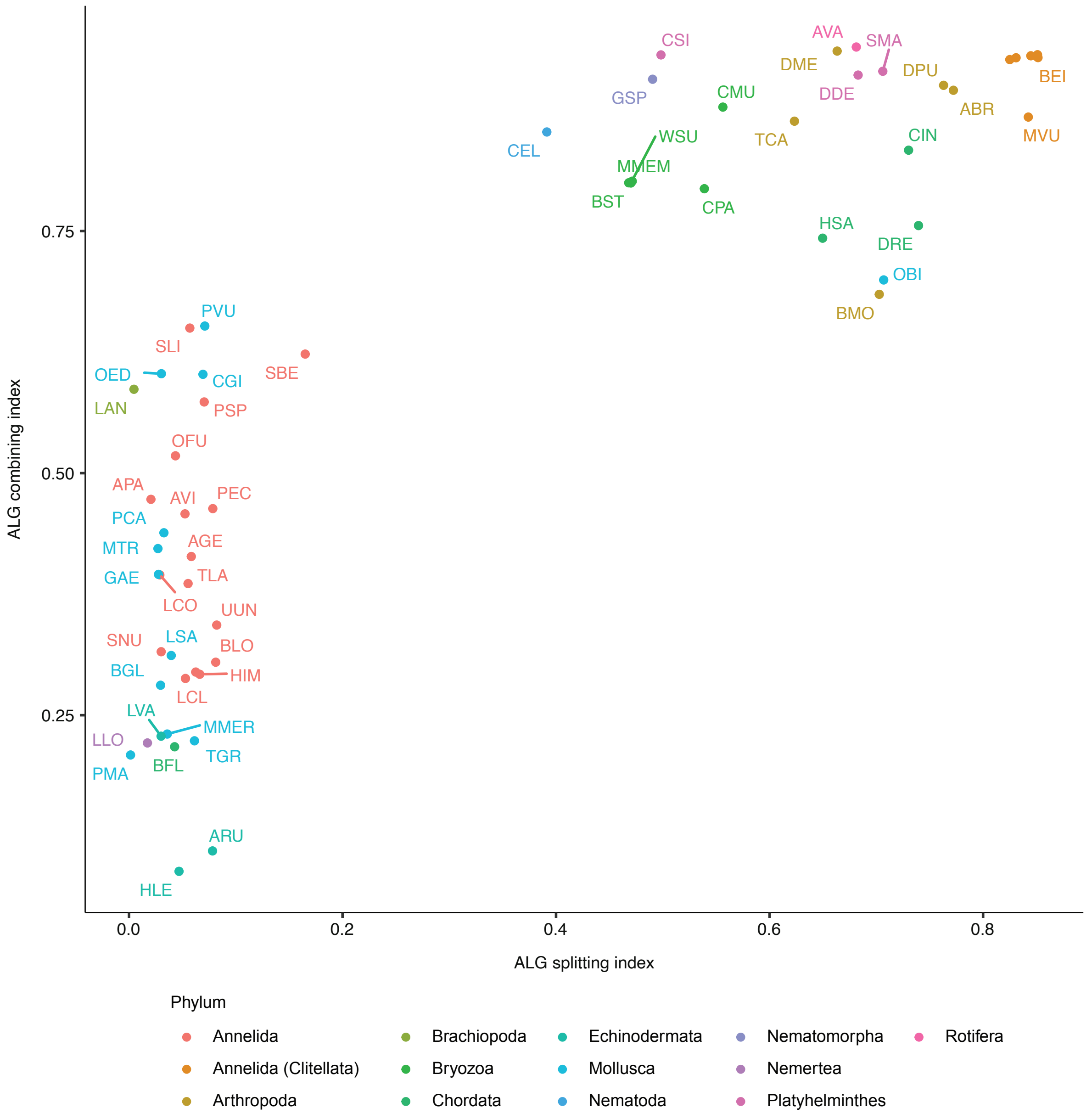


**Fig. S16. ALG splitting and ALG combining parameters for bilaterian species.** Plot is identical to that in main text Fig. 5B but with points colored by phylum. The plot supports the designation of two groups, one with high levels of both combining and splitting and one with low levels of splitting and varied levels of combining. Species abbreviations are in table S7.

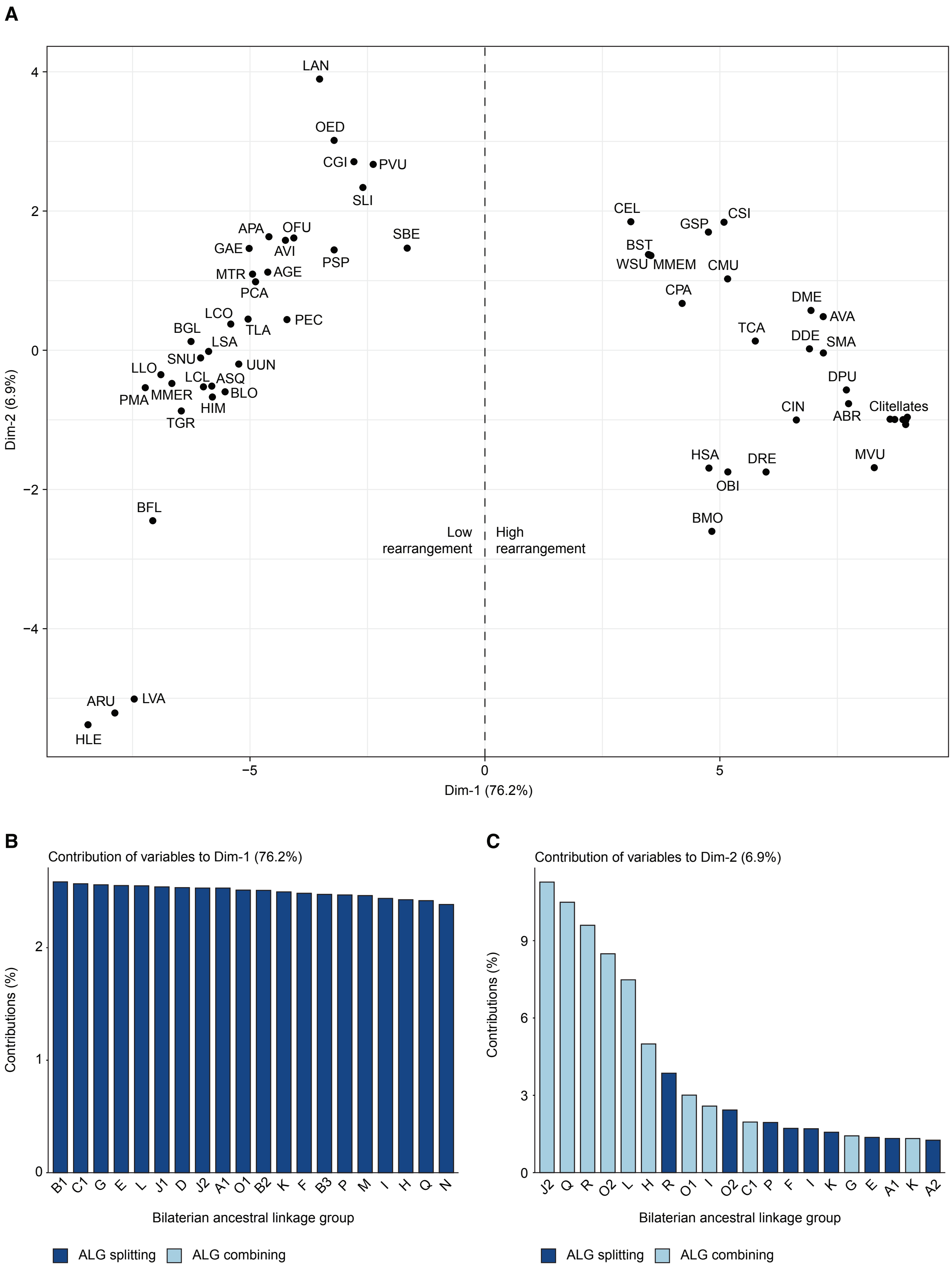

A
B
C
Low rearrangement
High rearrangement
Dim-1 (76.2%)
Dim-2 (6.9%)
LAN
OED
CGI
PVU
SLI
SBE
APA
OFU
GAE
AVI
PSP
MTR
AGE
PCA
LCO
PEC
BGL
TLA
LSA
SNU
UUN
LLO
LCL
ASQ
PMA
MMER
BLO
HIM
TGR
BFL
ARU
LVA
HLE
CEL
CSI
GSP
BST
WSU
MMEM
CMU
CPA
DME
AVA
TCA
DDE
SMA
DPU
CIN
ABR
Clitellates
HSA
DRE
MVU
OBI
BMO
Contribution of variables to Dim-1 (76.2%)
Contribution of variables to Dim-2 (6.9%)
Contributions (%)
Bilaterian ancestral linkage group
B1 C1 G E L J1 D J2 A1 O1 B2 K F B3 P M I H Q N
J2 Q R O2 L H R O1 I O2 C1 P F I K G E A1 K A2
ALG splitting
ALG combining

**Fig. S17. Principal component analysis (PCA) of rearrangement indices.** (**A**) PCA linear dimensionality reduction plot of ALG splitting indices and ALG combining indices for each individual species. The plot clearly separates low rearrangement species (left) from high rearrangement species (right) (**B**) Contribution of top 20 variables to Dim-1. ALG splitting variables and ALG combining variables are dark and light blue, respectively. ALG splitting is the major driver of variation in Dim-1. (**C**) Contribution of top 20 variables to Dim-2. ALG splitting variables and ALG combining variables are dark and light blue, respectively. ALG combining is the major driver of variation in Dim-2.

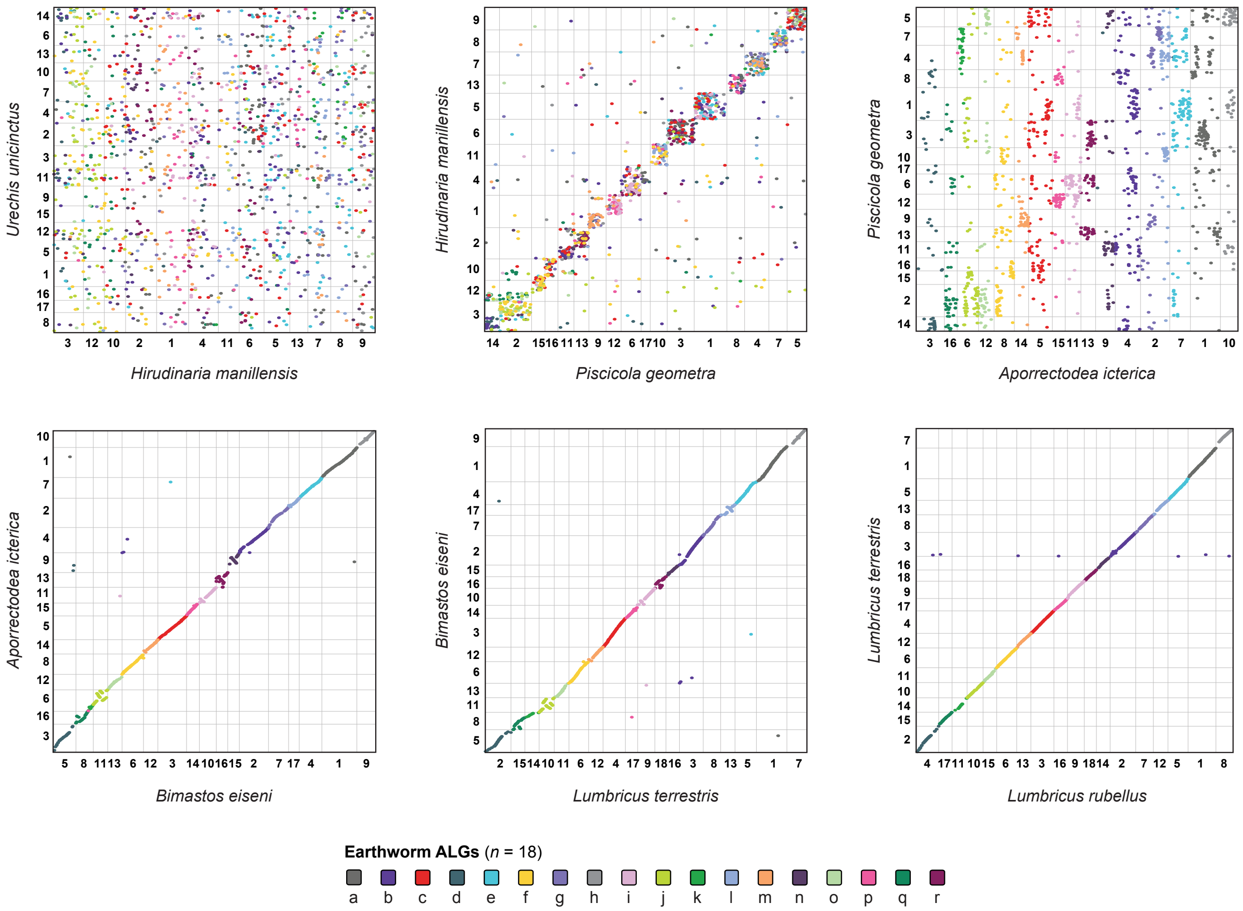


**Fig. S18. Oxford dot plots for clitellate synteny using earthworm ALGs.** Plots correspond to the ideogram plots in main text Fig. 6. Plots are colored by earthworm ALGs. Earthworm ALGs are highly conserved within earthworms, partially conserved between earthworms and leeches, and not conserved outside of clitellates.

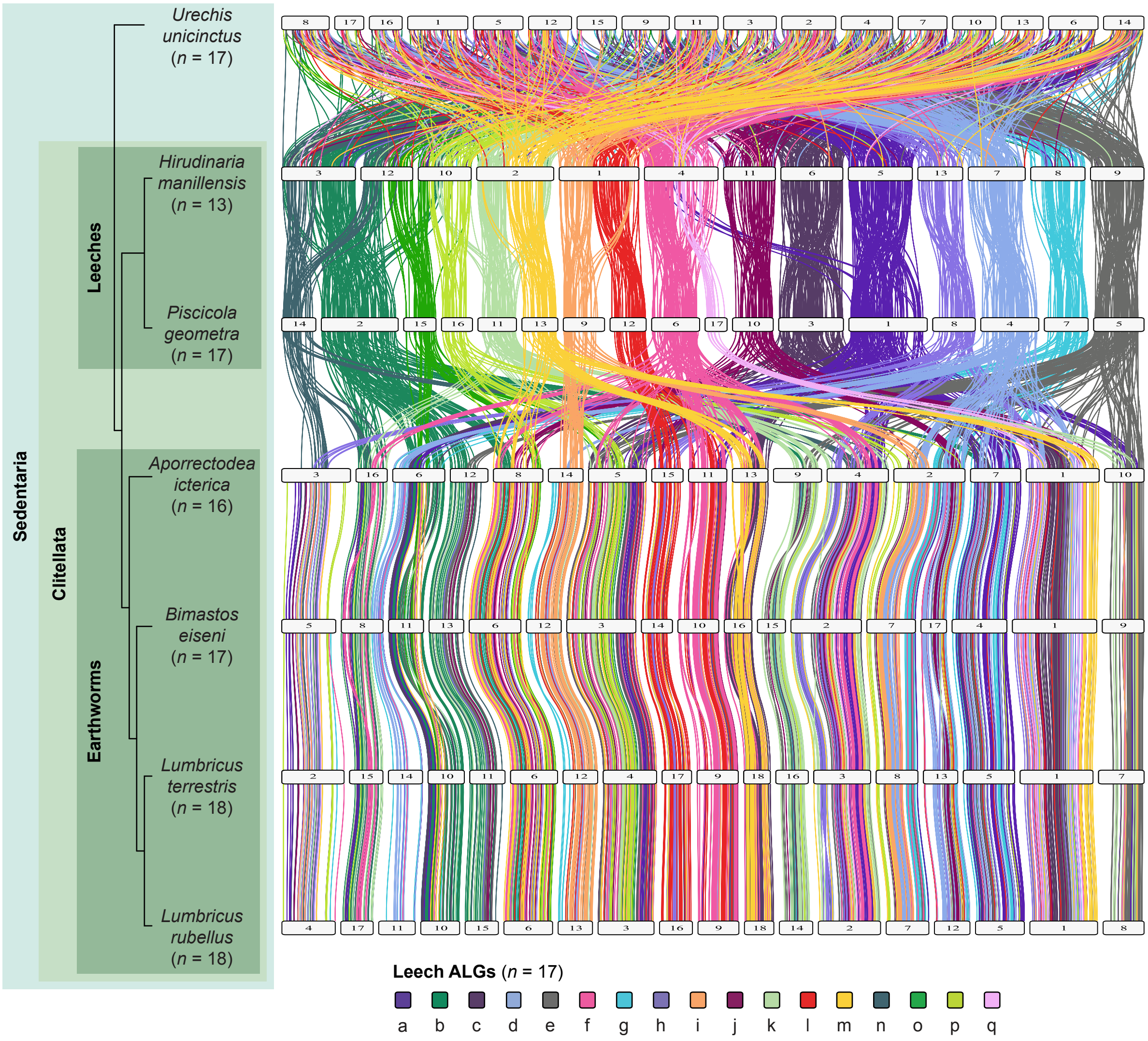


**Fig. S19. Clitellate synteny using leech ALGs.** Ideogram plots of clitellate genomes colored by leech ALGs. Leech ALGs are defined by genes' position in *Piscicola geometra*. Leech ALGs are largely conserved within the two leeches, partially conserved between leeches and earthworms, and completely unconserved in non-clitellate species like *Urechis unicinctus.*

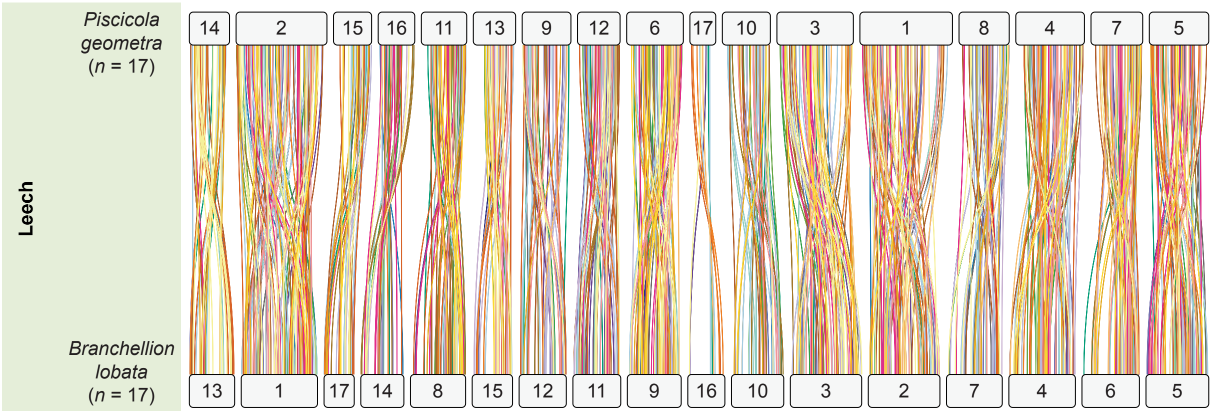


**Fig. S20. Macrosynteny of the leech *Branchellion lobata.*** *B. lobata* was excluded from the main analysis due to concerns over the low BUSCO score of its genome. Ideogram plot compares *B. lobata* to *Piscicola geometra*, which is in the same family (Piscicolidae). *B. lobata* and *P. geometra* chromosomes have a 1:1 relationship with no interchromosomal fusion or fission events.
